# CSF-1 maintains pathogenic but not homeostatic myeloid cells in the central nervous system during autoimmune neuroinflammation

**DOI:** 10.1101/2020.10.23.352534

**Authors:** Daniel Hwang, Maryam S. Seyedsadr, Larissa Lumi Watanabe Ishikawa, Alexandra Boehm, Ziver Sahin, Giacomo Casella, Soohwa Jang, Michael V. Gonzalez, James P. Garifallou, Hakon Hakonarson, Weifeng Zhang, Dan Xiao, Abdolmohamad Rostami, Guang-Xian Zhang, Bogoljub Ciric

## Abstract

The receptor for colony stimulating factor 1 (CSF-1R) is important for the survival and function of myeloid cells that mediate pathology during experimental autoimmune encephalomyelitis (EAE), an animal model of multiple sclerosis (MS). CSF-1 and IL-34, the ligands of CSF-1R, have similar bioactivities but distinct tissue and context-dependent expression patterns, suggesting that they have different roles. This could be the case in EAE, given that CSF-1 expression is upregulated in the CNS, while IL-34 remains constitutively expressed. We found that targeting CSF-1 with neutralizing mAb halted ongoing EAE, with efficacy superior to CSF-1R inhibitor BLZ945, whereas IL-34 neutralization had no effect, suggesting that pathogenic myeloid cells were maintained by CSF-1, not IL-34. Both anti-CSF-1- and BLZ945-treatment greatly reduced numbers of monocyte-derived cells and microglia in the CNS. However, anti-CSF-1 selectively depleted inflammatory microglia and monocytes in inflamed CNS areas, whereas BLZ945 depleted virtually all myeloid cells, including quiescent microglia, throughout the CNS. Anti-CSF-1 treatments reduced the size of demyelinated lesions, and microglial activation in the grey matter. Lastly, we found that bone marrow-derived immune cells were the major mediators of CSF-1R-dependent pathology, while microglia played a lesser role. Our findings suggest that targeting CSF-1 could be effective in ameliorating MS pathology, while preserving the homeostatic functions of myeloid cells, thereby minimizing risks associated with total ablation of CSF-1R-dependent cells.

**Significance Statement:** Multiple sclerosis (MS) and its animal model, experimental autoimmune encephalomyelitis (EAE), are autoimmune diseases characterized by accumulation of myeloid immune cells into the central nervous system (CNS). Both harmful and beneficial myeloid cells are present in EAE/MS, and a goal of MS therapy is to preferentially remove harmful myeloid cells. The receptor for CSF-1 (CSF-1R) is found on myeloid cells and it is important for their survival. CSF-1R can bind two ligands, CSF-1 and IL-34, but is unknown whether their functions in EAE/MS differ. We found that blocking CSF-1 depleted only harmful myeloid cells in the CNS and suppressed EAE, whereas blocking IL-34 had no effect. Thus, we propose that blocking CSF-1 could be a novel therapy for MS.

## INTRODUCTION

Multiple sclerosis (MS) is an autoimmune disease characterized by accumulation of immune cells in inflamed areas of the central nervous system (CNS), which eventually become demyelinated lesions [2]. Myeloid cells account for up to 85% of immune cells in active MS lesions [3–5], suggesting that they are the major mediators of pathology in MS. In support of this notion, studies in experimental autoimmune encephalomyelitis (EAE), an animal model of MS, have demonstrated the essential role of myeloid cells in EAE pathology, as interventions that affect them, in particular monocytes and conventional dendritic cells (cDCs), ameliorate or abrogate EAE [6–8]. However, despite evidence on the importance of myeloid cells in CNS autoimmunity, they have not been specifically targeted for MS therapy. This provides an opportunity for devising therapeutic approaches that target myeloid cells relevant to MS pathology.

Receptor for colony stimulating factor 1 (CSF-1R) is a cell-surface receptor tyrosine kinase that binds two ligands, CSF-1 (a.k.a. M-CSF) and IL-34 [9]. CSF-1R signaling facilitates survival and proliferation of myeloid cells, with either CSF-1 or IL-34 predominantly controlling the population size of myeloid cells in various organs and tissues [9–11]. CSF-1R is expressed by microglia, monocytes and monocyte-derived cells, which comprise the bulk of myeloid cells in the CNS during MS and EAE [10]. CSF-1R function is not intrinsically pro-inflammatory, as CSF-1R signaling in steady state induces a regulatory/homeostatic phenotype in macrophages, and a resting/quiescent phenotype in microglia [1, 12–26]. However, in inflammation, CSF-1R function could indirectly be pro-inflammatory, by simply perpetuating the survival and expansion of inflammatory myeloid cells. A recent study found an increase in both CSF-1R and CSF-1 in and around demyelinating lesions in cortical white matter of patients with progressive MS, and elevated CSF-1 in patients’ cerebrospinal fluid, whereas IL-34 was not increased, suggesting that CSF-1 drives the deleterious role of CSF-1R in MS [27]. In the progressive EAE model, CSF-1R has been identified as a key regulator of the inflammatory response in the CNS, with CSF-1R and CSF-1 expression levels correlating with disease progression. Inhibition of CSF-1R with small molecule inhibitor reduced the expression of pro-inflammatory genes and restored expression of homeostatic genes in microglia of mice with EAE [27].

It has been shown that direct inhibition of CSF-1R kinase activity with small molecule inhibitors suppresses EAE pathology [28, 29], but the effects of CSF-1R inhibition on particular cell subsets remain poorly understood. Different methods for reducing CSF-1R signaling, such as by antibodies (Ab) against the receptor or its individual ligands, have not been compared with small molecule inhibitors. The principal difference between these blocking methods is that small molecule inhibitors readily penetrate the entire CNS [1], whereas Abs have limited access, mainly to inflamed areas where the blood-brain barrier (BBB) has been compromised [10, 30]. Further, inhibitors of CSF-1R kinase activity could also affect other kinases [31], causing unwanted effects, especially over their prolonged use for therapy. In addition, direct inhibition of CSF-1R indiscriminately affects its systemic functions in their entirety, whereas individual neutralization of CSF-1 and IL-34 would have more nuanced effects, by preserving functions of the ligand that has not been targeted. These differences could lead to distinct outcomes in therapy of autoimmune neuroinflammation, given that the types and numbers of myeloid cells affected by various means of CSF-1R inhibition can substantially differ. In addition, any indirect effects of CSF-1R inhibition on cells that do not express CSF-1R also remain largely uncharacterized.

Our data and other studies show that small molecule inhibitors of CSF-1R cause a profound depletion of microglia [1, 10], which may be an important drawback in their use for therapy, as microglia play important roles in CNS homeostasis [23]. It has also been shown that neurons can express CSF-1R during excitotoxic injury, contributing to their survival [23]. Notably, it remains unknown whether neurons express CSF-1R in EAE and MS, and if its expression would be of significance, but, if so, restricting inhibition of CSF-1R signaling to inflamed areas of the CNS could be beneficial for neuronal survival. Thus, the ideal MS therapy would preferentially block CSF-1R signaling only in inflamed regions, affecting only inflammatory myeloid cells in them, while sparing cells in the rest of the CNS, such as quiescent microglia. This may be accomplished by targeting CSF-1R ligands, CSF-1 and IL-34, which show spatial- and context-dependent differences in expression [32]. Notably, although some minor differences have been described [33], CSF-1 and IL-34 have similar functions at the cellular level [34], suggesting that the differences found in animals lacking CSF-1 or IL-34 are primarily due to differential expression patterns [35]. IL-34 also binds to a second receptor, PTP-ζ [36], which has functions in oligodendrocyte development and homeostasis [37]. CSF-1 is systemically the dominant CSF-1R ligand, with its serum concentrations approximately ten times greater than that of IL-34 [15, 38–43]. Importantly, CSF-1 is not highly expressed in the CNS, but its expression can be upregulated by inflammation or injury [44], facilitating expansion of myeloid cells at the site of inflammation. In contrast to more widespread CSF-1 expression, IL-34 is primarily and constitutively expressed in the CNS and skin [45, 46]. In steady state, IL-34 maintains survival of tissue-resident myeloid cells in the skin and CNS, as the primary effect of IL-34 knockout is lack of Langerhan’s cells and microglia, respectively [32]. IL-34 is the predominant CSF-1R ligand in the CNS, accounting for 70% of total CSF-1R signaling in healthy brain [23].

In the present study, we sought to understand how CSF-1R inhibition affects immune cells in the CNS of mice with EAE, and to determine how the effects of blocking CSF-1 and IL-34 may differ from blocking CSF-1R. We found that treatment with CSF-1R inhibitor BLZ945 suppresses EAE when given both prophylactically and therapeutically. Treatment efficacy correlated with a dramatic reduction in the numbers of monocytes, monocyte-derived dendritic cells (moDCs), and microglia, suggesting that loss of one or more of these cell types is responsible for EAE suppression. We also found that blockade of CSF-1, but not of IL-34, with Ab suppressed EAE. Importantly, anti-CSF-1 treatment preferentially depleted inflammatory myeloid cells, whereas quiescent microglia were preserved. These findings suggest that blocking CSF-1, rather than CSF-1R, may be a suprerior therapeutic strategy for alleviating pathology in MS.

## RESULTS

### Blocking CSF-1R or CSF-1, but not IL-34, suppresses EAE

To determine the roles of CSF-1R, CSF-1, and IL-34 in EAE, we blocked them with either a small-molecule inhibitor or neutralizing monoclonal Abs (MAbs). We blocked CSF-1R kinase function with BLZ945, a CNS-penetrant small-molecule inhibitor that, among other effects, potently depletes microglia within several days [1]. Treatment with BLZ945 delayed the onset of EAE for 6-15 days when given prophylactically (Fig. 1A) and initially suppressed disease severity but could not halt disease progression despite continuous treatment. We also blocked CSF-1R activity with neutralizing MAb, which was less effective than BLZ945 in delaying EAE onset, and reducing disease severity (Fig. 1B). Surprisingly, anti-CSF-1R MAb (clone AFS98) failed to bind to microglia, as determined by flow cytometry, whereas it bound to monocytes (Supplemental Fig. 1A). It is unclear why this MAb did not bind to microglia, and how that may have impacted its effect in EAE.

**Figure 1:**
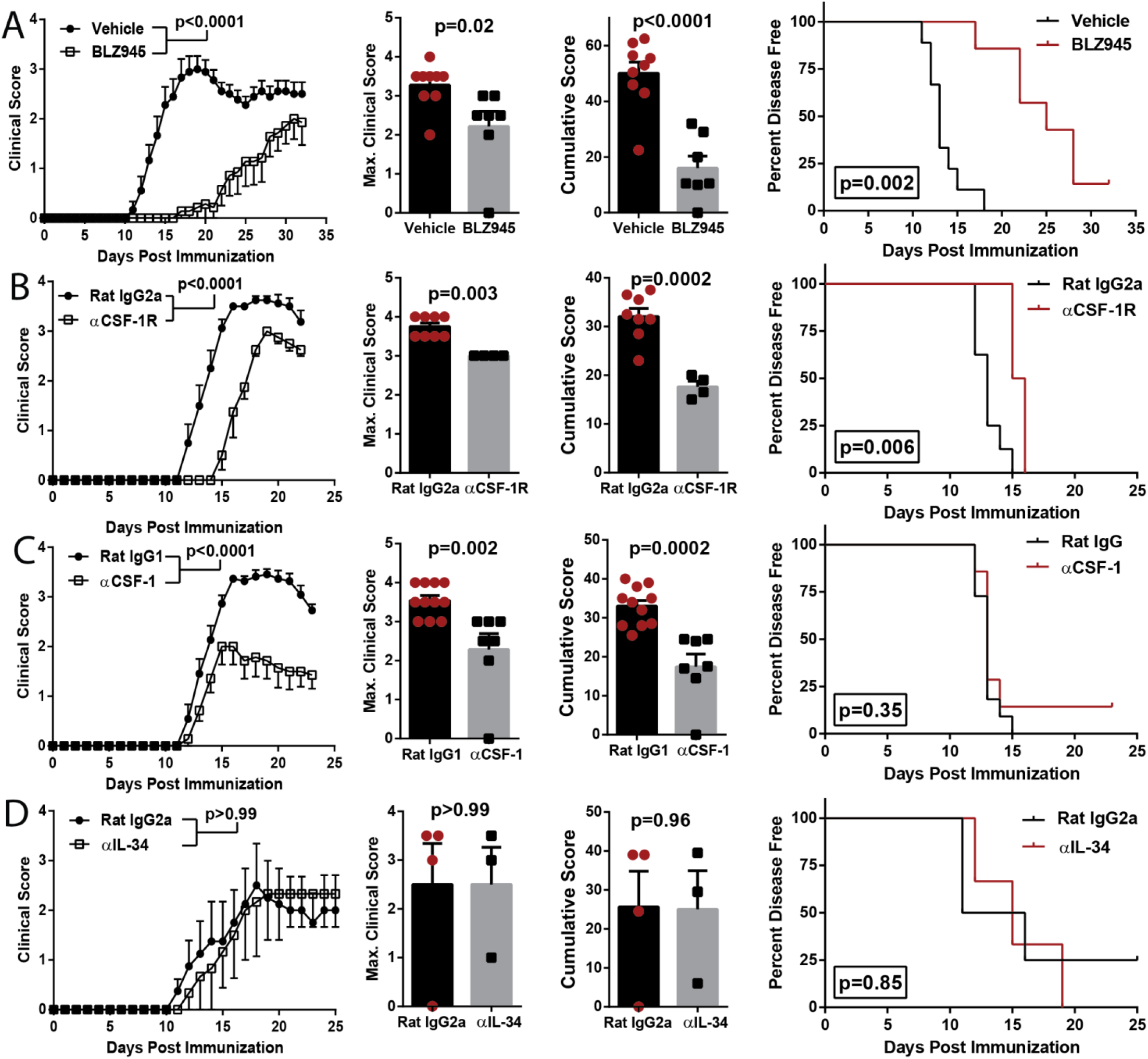
Blocking CSF-1R activity suppresses EAE. **A-D)** C57BL/6J mice were immunized with MOG35-55 for EAE induction. Clinical course, maximum and cumulative clinical scores, and Kaplan-Meier plots depicting percent of disease-free animals over time are shown. Significance for clinical course data was calculated by two-way repeated measures ANOVA. Significance for maximum and cumulative clinical scores was calculated by Student’s *t* test. Error bars are S.E.M. Significance for Kaplan-Meier plots was calculated by comparing disease-free curves with the log-rank (Mantel-Cox) test. **A)** EAE animals treated orally with BLZ945 (n=9; 4 mg/day [1]) or vehicle (n=7; 20% Captisol) daily, starting from day of immunization. Data were compiled from two independent experiments. **B)** Treatment with anti-CSF-1R MAb (n=4) or control rat IgG2a (n=8). MAbs were i.p. injected every other day (400 µg per dose). **C)** Treatment with anti-CSF-1 MAb (n=7) or control rat IgG1 (n=11). MAbs were i.p. injected every other day (200 µg per dose). Data were compiled from two independent experiments. **D)** Treatment with anti-IL-34 MAb (n=3) or control rat IgG2a MAb (n=4). MAbs were i.p. injected every other day (100 µg per dose).

We then tested how blocking either CSF-1 or IL-34 with MAbs would affect EAE. In contrast to blocking CSF-1R, blocking of CSF-1 did not delay onset of disease (Fig. 1C), but did continuously suppress disease severity over the course of treatment, including for over 40 days (Fig. 3B). Anti-IL-34 treatment did not suppress EAE (Fig. 1D). We tested whether our treatments with the MAbs, which were rat IgGs, induced an anti-rat IgG response in mice with EAE. Anti-CSF1, and control rat IgG2a isotype MAb induced similar low titers of anti-rat IgG, whereas anti-CSF-1R and anti-IL-34 MAbs (both rat IgG2a) induced notably higher anti-rat IgG response (Supplemental Fig. 1B), indicating that anti-rat IgG responses in treated mice may have reduced the neutralizing effects of injected MAbs in the case of anti-CSF-1R and anti-IL-34 MAbs. To minimize the development of anti-rat IgG responses, we treated mice with anti-IL-34 MAb immediately after the onset of clinical disease; however, this treatment had no impact on disease (Supplemental Fig. 1C). We also tested whether administration of recombinant CSF-1 could exacerbate EAE pathology, but i.p. injections of CSF-1 did not worsen disease (Supplemental Fig. 1D). Overall, these data show that blocking CSF-1R signaling attenuates EAE, and that CSF-1, but not IL-34, is the relevant CSF-1R ligand for EAE pathology.

### Inhibition of CSF-1R signaling depletes myeloid APCs in the CNS during EAE

We characterized how prophylactic treatment with BLZ945 influenced CNS inflammation at the peak of EAE. BLZ945-treated animals had ∼90% reduced numbers of CD45^+^ cells in the CNS compared to vehicle-treated animals (Fig. 2A). All major types of immune cells were reduced in number, but CD11b^+^ cells were most impacted, including profound depletion of CD45^Low^CD11b^+^Tmem119^+^CX3CR1^Hi^ microglia and CD45^Hi^CD11b^+^CD11c^+^ myeloid DCs, including CD45^Hi^CD11b^+^CD11c^+^Ly6G^Low/-^Ly6C^Hi^MHCII^Hi^ moDCs (Supplemental Fig. 2). We clustered flow cytometry data by *t*-stochastic neighbor embedding (*t*-SNE) and consistent with manual gating, the frequency of microglia and moDCs/macrophages was dramatically reduced (Fig. 2B-E). In contrast, the frequency of neutrophils was increased; likely reflecting that they do not express CSF-1R [47], and are therefore not impacted by CSF-1R inhibition. Interestingly, the frequency of undifferentiated monocytes (CD45^Hi^CD11b^+^Ly6C^Hi^CD11c^-^MHCII^-^) increased, suggesting that CSF-1R may impact monocyte differentiation or their survival/proliferation after initial differentiation. Overall, BLZ945 treatment markedly reduced the frequency of CD11c^+^ myeloid antigen-presenting cells (APCs) expressing MHC class II, CD80 and CD86 (Fig. 2E,F), suggesting that CSF-1R signaling maintains sufficient numbers of APCs in the CNS to drive inflammation during EAE. Lastly, we found differences in cytokine production by CD4^+^ T cells from BLZ945-treated mice, including lower frequency of GM-CSF^+^ cells (Supplemental Fig. 2G).

**Figure 2:**
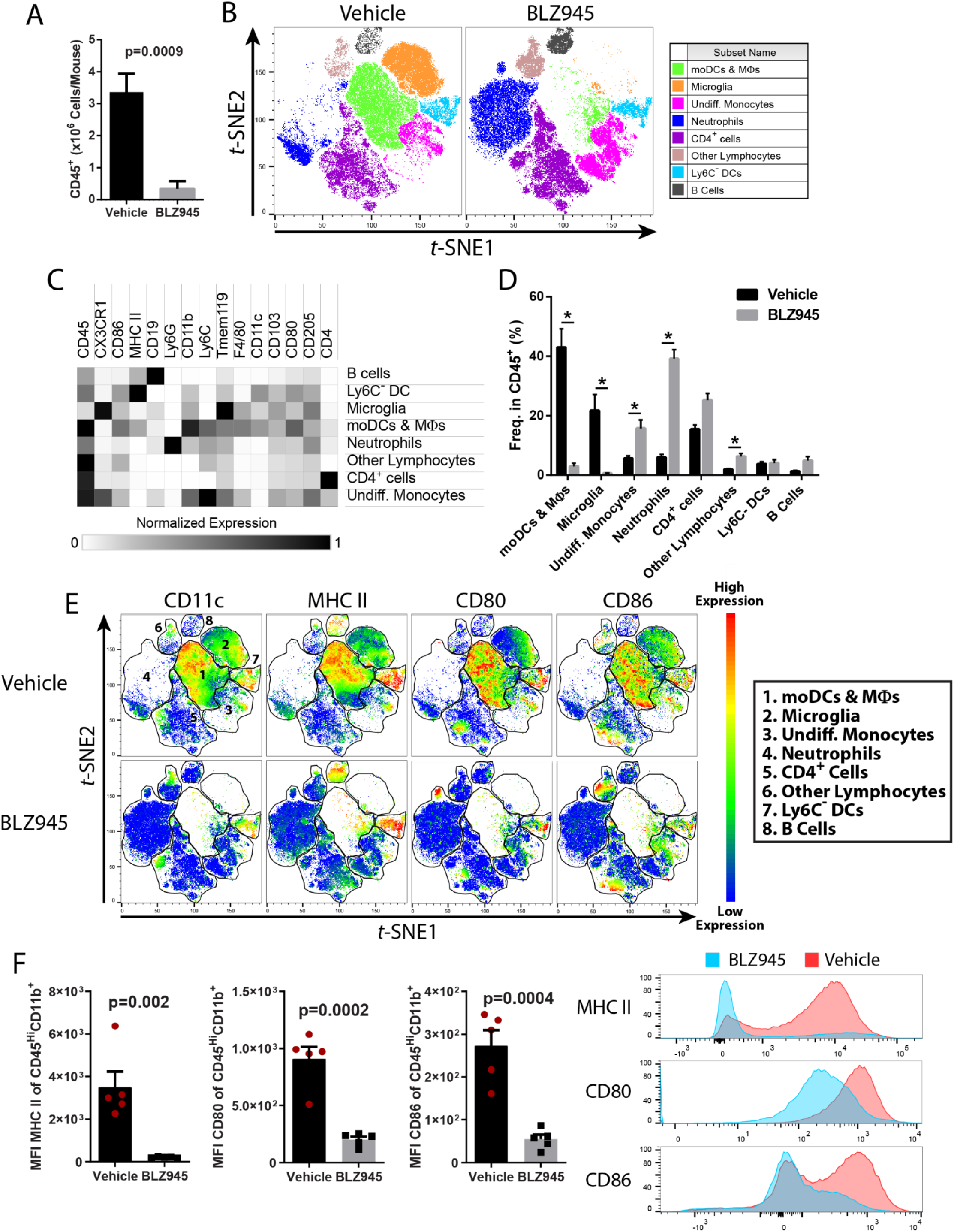
CSF-1R inhibition depletes myeloid APCs in the CNS of mice with EAE. C57BL/6J mice were immunized with MOG35-55 for EAE induction and treated orally with BLZ945 (4 mg/day) or vehicle control (20% captisol) daily, starting on the day of immunization. Mice were sacrificed on day 15 p.i., and brains and spinal cords were pooled for cell isolation. **A)** Numbers of CNS CD45^+^ cells (n=7/group, combined from two independent experiments). **B)** *t*-SNE plot depicting clustering of CD45^+^ cells (n=5 mice per group). moDCs and macrophages were defined by their expression of CD11b, CD11c, MHC II, CD80 and CD86. Microglia were defined as CD45^+^CD11b^+^Tmem119^+^CX3CR1^Hi^ cells. Undifferentiated monocytes were defined as CD45^Hi^Ly6C^Hi^CD11c^-^ cells that were overall MHC II^Lo/Neg^. Neutrophils were defined as CD45^+^CD11b^+^Ly6G^Hi^. CD4^+^ cells were defined as CD45^+^CD4^+^. Other lymphocytes were defined as CD45^Hi^SSC^Lo^. Ly6C^-^ DCs were defined as CD45^Hi^CD11b^+^CD11c^+^MHCII^Hi^Ly6C^-^ cells. B cells were defined as CD45^+^CD19^+^. **C)** Heatmap showing normalized expression of markers used to identify clusters. **D)** Quantification of clusters between vehicle and BLZ945-treated mice with EAE. **E)** Heat map of CD11c, MHC II, CD80 and CD86 expression among CD45^+^ cells. **F)** MFI of MHC II, CD80 and CD86 among CD45^Hi^CD11b^+^ myeloid cells. Representative histograms showing fluorescence intensity of MHC II, CD80 and CD86 between vehicle- and BLZ945-treated animals is also shown. Y-axis is frequency of cells as normalized to mode. Significance was calculated with Student’s *t* test. For D),* indicates a p-value < 0.02. Error bars are S.E.M.

We next tested BLZ945-treatment during ongoing EAE, as that scenario is the most relevant to MS therapy. Therapeutic treatment with 200mg/kg/day BLZ945 suppressed clinical EAE, with better efficacy being observed at 300 mg/kg/day (Supplemental Fig 3A-B). To determine the acute effects of CSF-1R inhibition on immune cells in the CNS, we focused our analysis on mice treated with BLZ945 for 6 days, starting at a clinical score of 2.0. BLZ945-treated mice had reduced numbers of CD45^+^ cells in the CNS, primarily due to fewer CD11b^+^ and CD11c^+^ cells, including microglia and CD45^Hi^CD11b^+^CD11c^+^ myeloid DCs (Supplemental Fig. 3C-F). Most immune cells depleted by BLZ945 treatment co-expressed CD11c, TNF, MHCII, CD80, and CD86, suggesting that inflammatory APCs are preferentially affected by CSF-1R inhibition (Supplemental Fig. 3G-I). Taken together, these data further indicate that inhibition of CSF-1R signaling suppresses EAE by reducing numbers of APCs in the CNS.

### Blocking CSF-1 depletes inflammatory myeloid APCs in the CNS without affecting quiescent microglia

We next tested the therapeutic efficacy of anti-CSF-1 MAb treatment. Treatment initiated after onset of disease suppressed clinical EAE (Fig. 3A), and the suppression was maintained up to 45 days after EAE induction, which was the longest period tested (Fig. 3B). Similar to BLZ945, anti-CSF-1 MAb suppressed disease even when treatment was initiated during its more advanced stage (Fig. 3C, D). Mice treated with anti-CSF-1 MAb had fewer CD45^+^ immune cells in the CNS, including CD11b^+^, CD11c^+^, and CD4^+^ cells (Fig. 3E, F). The treatment decreased frequency of CD11b^+^CD11c^+^ myeloid DCs in the CNS (Supplementarl Fig. 4A), and among CD11b^+^ myeloid cells, we observed reduced frequencies of CD11c^+^ microglia, moDCs, cDCs and other CD11c^+^ cells, indicating that DC populations were preferentially affected (Supplementary Fig. 4B). Anti-CSF-1 treatment reduced the numbers of microglia to those in naïve mice (Fig. 3G), without depleting the entirety of microglia, as was the case with BLZ945. Importantly, the reductions in microglia numbers in anti-CSF-1-treated mice were primarily due to loss of activated inflammatory microglia, which expressed MHC II and/or CD68 (Fig. 3H).

**Figure 3:**
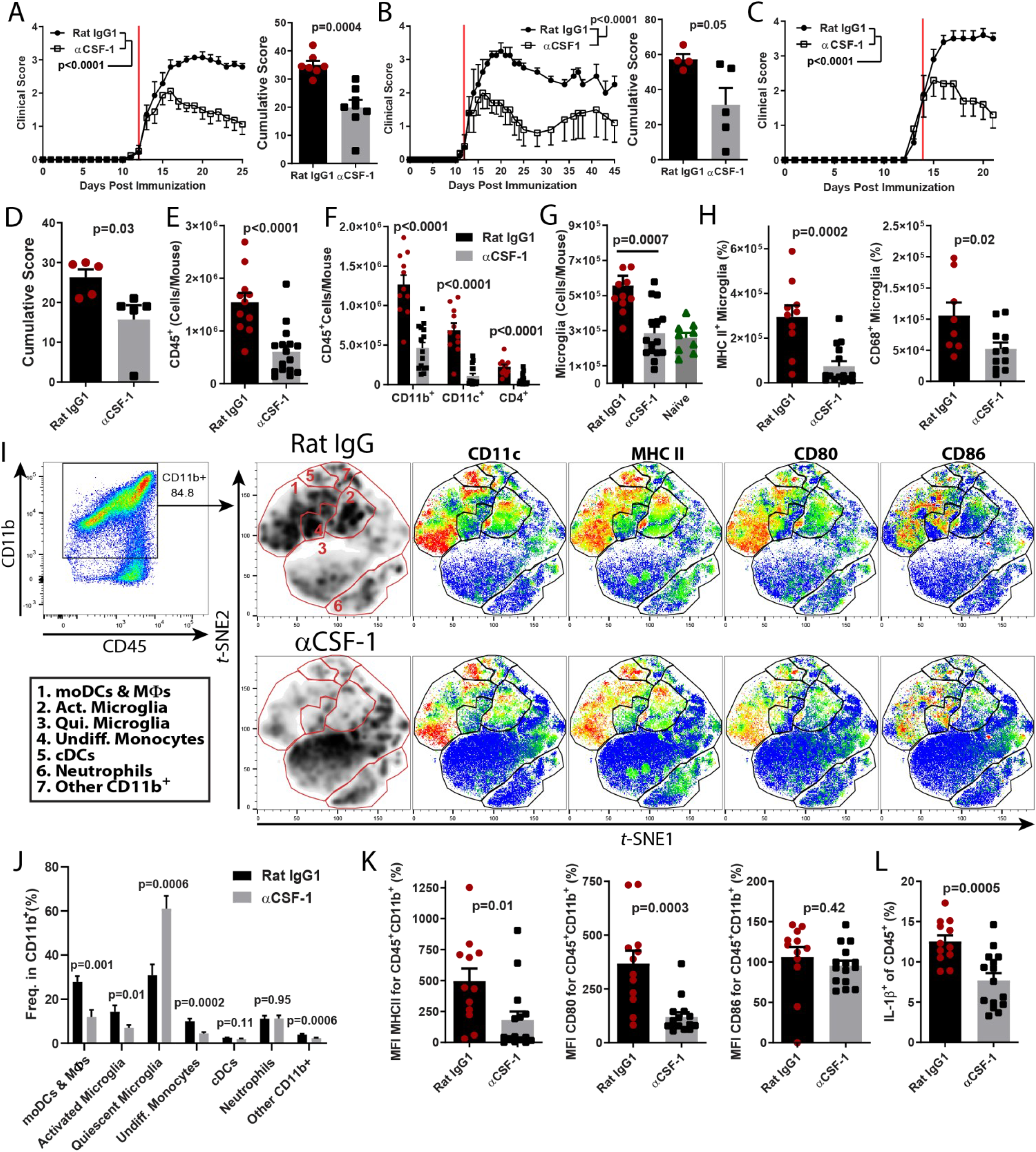
CSF-1 controls the population size of inflammatory myeloid cells in the CNS during EAE. **A)** Clinical course and cumulative score of mice with EAE treated with αCSF-1 MAb starting after disease onset (n=7 per group; compiled from 2 independent experiments). Red line denotes day that treatment was started. Mice were treated with 200 μg MAb per day. **B)** Clinical course and cumulative scores of mice treated long term with αCSF-1 MAb. Mice were treated daily until day 25, and then every other day for the duration of the experiment. **C-D)** Clinical course and cumulative score for mice treated with αCSF-1 MAb, starting at clinical score of 2.0 (n=4-5 per group). **E-L)** Characterization of immune cells from the CNS of control MAb- and αCSF-1 MAb-treated mice. **E)** Number of CD45^+^ cells. **F)** Numbers of CD11b^+^, CD11c^+^ and CD4^+^ cells. **G)** Numbers of CD45^+^CD11b^+^CX3CR1^Hi^Tmem119^+^ microglia in control-treated, αCSF-1 MAb-treated, and naïve mice. **H)** Numbers of MHC II^+^ and CD68^+^ microglia in control MAb- and αCSF-1 MAb-treated mice. **I)** *t-*SNE analysis of CD45^+^CD11b^+^ cells. moDCs and macrophages were defined by their expression of CD11b, CD11c, MHC II, CD80 and CD86. Activated microglia were defined as CD45^+^CD11b^+^Tmem119^+^CX3CR1^Hi^MHCII^+^CD68^+/-^ cells. Quiescent microglia were defined as CD45^+^CD11b^+^Tmem119^+^CX3CR1^Hi^MHCII^-^CD68^-^. Undifferentiated monocytes were defined as CD45^Hi^Ly6C^Hi^CD11c^-^ cells that were overall MHC II^Lo/Neg^. Neutrophils were defined as CD45^+^CD11b^+^Ly6G^Hi^. CD4^+^ cells were defined as CD45^+^CD4^+^. cDCs were defined as CD45^Hi^CD11b^+^CD11c^+^MHCII^Hi^Ly6C^-^CD26^+^ cells. Other CD11b^+^ cells expressed CD11c and CX3CR1 but did not express markers for antigen presentation. **J)** Quantification of clusters from I). **K)** Median fluorescence intensity of MHC II, CD80 and CD86 in CD45^+^CD11b^+^ cells. **L)** Frequency of IL-1β^+^ cells among CD45^+^ cells. Significance was calculated with unpaired Student’s *t* test. Error bars are S.E.M. Significance for clinical course was calculated by two-way ANOVA.

We then quantified how anti-CSF-1 treatment affected the composition of myeloid cells in the inflamed CNS (Fig. 3I,J). Anti-CSF-1 treatment reduced the frequencies of moDCs, macrophages, activated microglia and undifferentiated monocytes, but did not affect the frequency of neutrophils. This resulted in a large increase in the frequency of quiescent microglia among CD11b^+^ cells. Similar to BLZ945-treated mice, anti-CSF-1 treatment resulted in a decrease in MFI for MHC II and CD80, but not CD86 (Fig. 3K), suggesting that inflammatory APCs were preferentially impacted by anti-CSF-1 treatment. Consistent with this, anti-CSF-1-treated mice had a lower frequency of TNF^+^MHCII^+^ cells in their CNS, including moDCs (Supplemental Fig. 4C,D). An important pathogenic function of moDCs in EAE is production of IL-1β [48]. As expected, a reduced frequency of IL-1β-producing cells was also observed in the CNS of anti-CSF-1 treated EAE mice (Fig. 3L), but, notably, not in BLZ945-treated mice (Supplemental Fig. 3I). Together, these data suggest that blockade of CSF-1 preferentially depletes infiltrating and resident inflammatory myeloid cells, without affecting quiescent microglia.

### Anti-CSF1 treatment depletes myeloid cells in white matter lesions, but not microglia in grey matter

We further tested whether treatment with anti-CSF-1 MAb preferentially depletes inflammatory myeloid cells by examining their distribution in the CNS by microscopy. Anti-CSF-1 MAb treated mice with EAE had reduced white matter demyelination and smaller spinal cord lesions when compared to control mice (Fig. 4A-C). Immunostaining and analysis by confocal microscopy showed that mice treated with control isotype MAb had large numbers of Iba^+^ cells throughout the spinal cord (Fig. 4D-G). In contrast, mice treated with anti-CSF-1 MAb had notably fewer Iba^+^ cells and were similar to healthy naïve controls. BLZ945 treatment depleted virtually all Iba^+^ cells in both white and grey matter. Anti-CSF-1-treated mice also had reduced density of Iba1^+^ and Iba^+^CD68^+^ inflammatory myeloid cells both within regions infiltrated with immune cells and in immediately surrounding areas, but not further into the grey matter (Fig. 4E,F). Treatment with anti-CSF-1 reduced microglial proliferation/activation in the grey matter, reducing numbers of grey matter microglia to levels found in naïve mice (Fig. 4G). Together, these data indicate that anti-CSF-1 treatment primarily targets inflammatory myeloid cells in CNS lesions without depleting quiescent microglia in the grey matter. Further, the depletion of inflammatory myeloid cells in inflamed areas of the white matter precluded alterations to microglia in normally-appearing white and gray matter at sites distant from inflamed areas of the white matter [4, 49–51], indicating an overall reduction in CNS inflammation.

**Figure 4:**
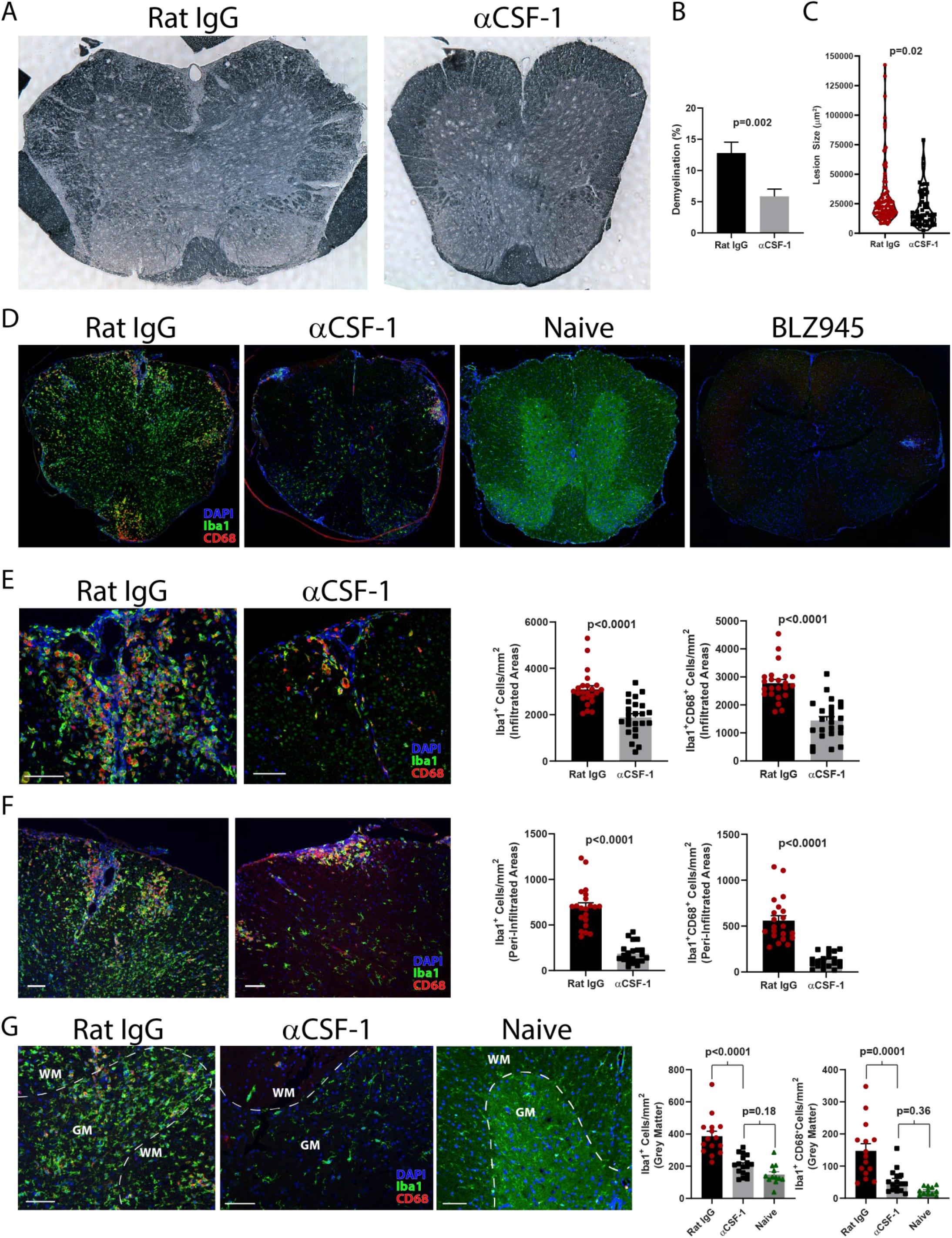
Anti-CSF1 treatment reduces lesion burden without depleting quiescent microglia. **A)** Sudan black stained spinal cord sections from rat IgG- and anti-CSF-1-treated mice with EAE sacrificed on day 17-19 p.i. (n=5 mice per group). Spinal cords were split into 4 pieces of equal length, and 1 section from each piece is included in the analysis. **B)** Degree of demyelination expressed as a percentage of demyelinated white matter. **C)** Violin plot showing demyelinating lesion size in rat IgG- and anti-CSF-1-treated mice. **D)** Confocal microscopy of spinal cord sections stained with DAPI and Abs against Iba1 and CD68 from rat IgG-, anti-CSF-1-, and BLZ945-treated EAE mice, and naïve mice. **E)** Representative images and quantification of cell density of Iba1^+^ and Iba1^+^CD68^+^ cells within lesions infiltrated with immune cells from rat IgG- and anti-CSF-1-treated mice. **F)** Representative images and quantification of cell density of Iba1^+^ and Iba1^+^CD68^+^ cells within a 100 µm-wide area surrounding lesions infiltrated with immune cells. **G)** Representative images and quantification of cell density of Iba1^+^ and Iba1^+^CD68^+^ cells in grey matter of rat IgG- or anti-CSF-1-treated and naïve mice. All scale bars are 100 µm. For E) and F), individual lesions from 5 mice per group are analyzed. Significance was calculated with unpaired Student’s *t* test. Error bars are S.E.M. Significance for comparisons between more than two groups was calculated by one-way ANOVA.

### CSF-1R inhibition depletes myeloid DCs and monocytes in peripheral lymphoid compartments

Treatment with BLZ945 delayed onset of disease, whereas anti-CSF-1 treatment did not. This difference could be due to diminished priming of encephalitogenic T cell responses in peripheral lymphoid organs of BLZ945-treated mice, resulting in failure to initiate disease in the CNS. To test this possibility, we treated immunized mice with either BLZ945 or anti-CSF-1 MAb and sacrificed them during the priming phase of EAE, on day 8 p.i. We quantified the immune cells in blood, draining lymph nodes (dLN) and spleen, and found no difference in overall numbers of CD45^+^ cells in any tissues examined from BLZ945- or anti-CSF-1-treated mice compared to control animals (Supplemental Fig. 5). We did, however, observe a decrease in the numbers of CD11b^+^ cells in all tissues examined from BLZ945-treated mice, but not from anti-CSF-1-treated mice (Supplemental Fig. 5A-C). We noted a decrease in CD11b^+^CD11c^+^ cells in some peripheral lymphoid organs in both BLZ945- and anti-CSF-1-treated mice, including a decrease in moDCs. Notably, numbers of monocytes were reduced in all examined tissues from BLZ945-treated mice, but not from anti-CSF-1-treated mice (Supplemental Fig 6A-F).

We tested whether reductions in myeloid DCs in the spleen and dLNs would diminish MOG_35-55_-specific T cells responses, but did not find reduced proliferation of cells from either BLZ945- or anti-CSF-1-treated animals when compared to control animals (Supplemental Fig. 6G,H). We also measured antigen-specific proliferation at day 16 p.i. and found a reduction in proliferation of splenocytes from BLZ945-treated mice, but not of cells from dLNs. The reduction was likely due to fewer APCs, rather than to intrinsic differences in APC function, as co-culture of equal numbers of CD11c^+^ cells purified from spleens of vehicle- or BLZ945-treated mice with CD4^+^ T cells from 2D2 mice elicited similar levels of proliferation (Supplemental Fig. 6I,J). These data show that myeloid DCs are impacted by CSF-1R inhibition, but that this only modestly affects the development of myelin antigen-specific responses. Thus, delayed onset of disease in BLZ945-treated animals is likely due to factors other than impaired development of MOG_35-55_-specific T cells responses.

### CSF-1R signaling promotes survival/proliferation of BM-derived moDCs, but not their APC function

We tested how CSF-1R signaling influences the numbers of DCs by generating BMDCs with GM-CSF and IL-4 [52], and neutralizing CSF-1 and CSF-1R with MAbs. Cultures with either anti-CSF-1 or anti-CSF-1R MAbs contained fewer CD11c^+^MHCII^Hi^ DCs (Fig. 5A). This was correlated with a decreased ratio of live/dead cells after LPS treatment (Fig. 5B), suggesting that survival of BMDCs was negatively impacted by the absence of CSF-1R signaling. CSF-1R signaling was not important for development of APC function of BMDCs as there was only a small reduction in the frequency of CD11c^+^MHCII^+^ among live CD11b^+^ cells (Fig. 5C,D) and co-culture with 2D2 CD4^+^ T cells revealed no differences in eliciting the proliferation of 2D2 T cells (Fig. 5E). To confirm that these findings are applicable to monocyte-derived BMDCs, we purified CD11b^+^Ly6G^-^Ly6C^Hi^ monocytes from the BM of CD45.1^+^ mice, mixed them with total BM cells from CD45.2^+^ mice, and then blocked CSF-1R signaling during their development into BMDCs. Consistent with total BM cultures, blockade of CSF-1R signaling did not affect the frequency of CD11c^+^MHCII^Hi^CD45.1^+^ monocyte-derived cells (Fig. 5F,G), but caused a ∼75% reduction in their numbers (Fig. 5H). Together, these data indicate that CSF-1R signaling promotes the survival of moDCs, rather than their differentiation and APC function, which is consistent with the role of CSF-1R signaling in maintaining myeloid cell populations [9–11].

**Figure 5:**
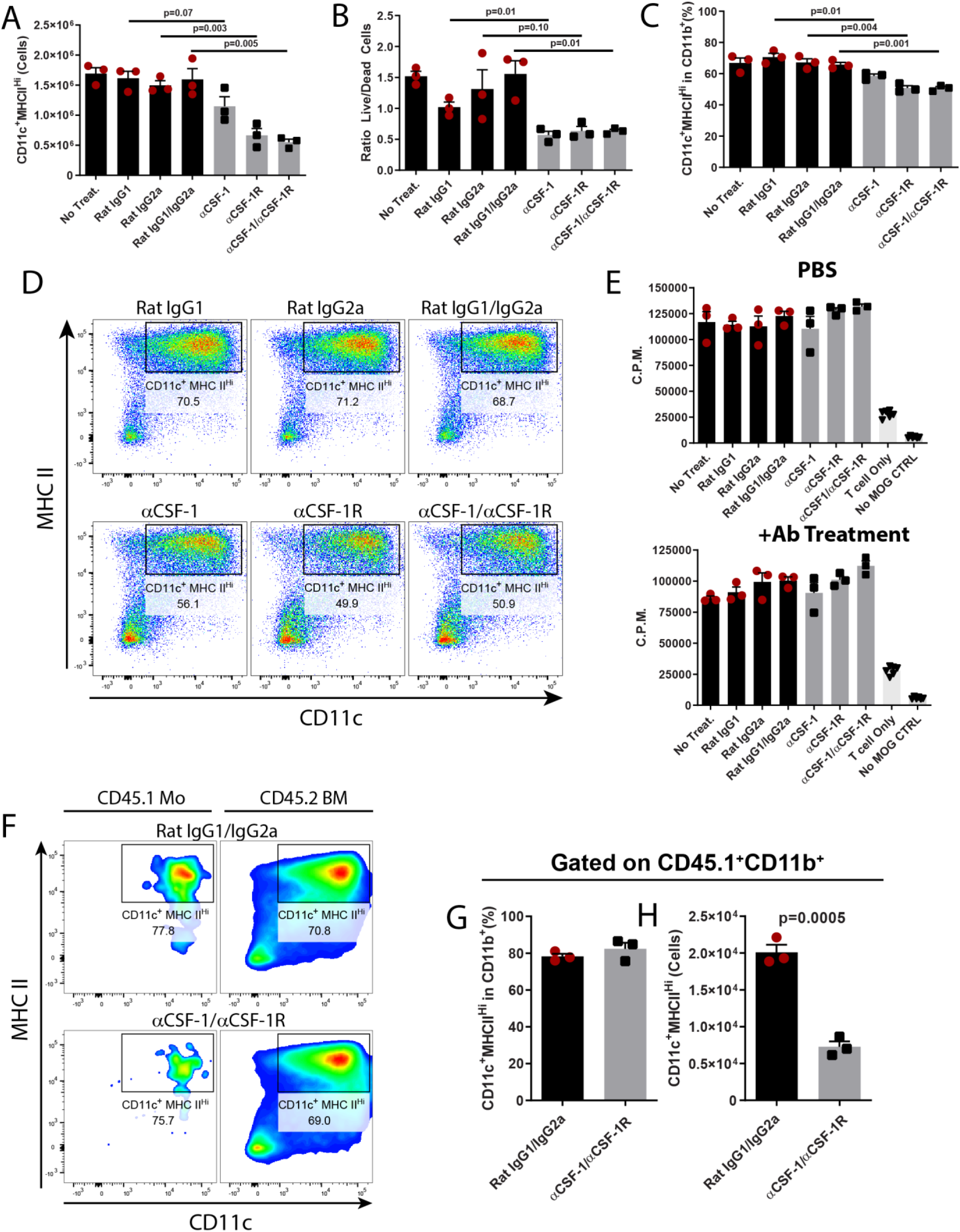
CSF-1R signaling promotes survival and proliferation of moDCs, but not their APC function. BM cells were cultured in media supplemented with GM-CSF and IL-4 for 7 days in the presence of either control MAbs, or anti- CSF1 and anti-CSF-1R MAbs. BMDCs were then matured by stimulation with LPS for 24 h in the presence of the MAbs. **A)** Number of CD11c^+^MHCII^Hi^ cells in culture after GM-CSF + IL-4 treatment for 7 days. **B)** Ratio of Live/Dead cells after 24 h LPS treatment. **C-D)** Frequency of CD11c^+^MHCII^Hi^ cells after GM-CSF + IL-4 treatment for 7 days. **E)** Co-culture of LPS-matured BMDCs with CD4^+^ T cells from 2D2 mice and MOG35-55 peptide (25 μg/mL). Co-cultures either did, or did not contain control and neutralizing MAbs against CSF-1 and CSF-1R. Proliferation was measured using [^3^H]Thymidine incorporation. C.P.M. = counts per minute. **F)** Monocytes were purified from the BM of CD45.1 mice and mixed with total BM of CD45.2 mice, then cultured as described above with control or anti-CSF1/anti-CSF-1R MAbs. Flow cytometry depicting CD11c and MHC II expression in CD45.1^+^CD11b^+^ and CD45.2^+^CD11b^+^ cells is shown. **G)** Frequency, and **H)** number of CD11c^+^MHC II^Hi^ cells among CD45.1^+^CD11b^+^ cells. Technical replicates for 1 of 2 independent experiments with similar results are shown. Statistical significance was calculated using two-way unpaired *t* test. Error bars are S.E.M.

### Blocking CSF-1R signaling or CSF-1 reduces numbers of CCL2- and CCR2-expressing myeloid cells in the CNS during EAE

We observed that numbers of monocytes/moDCs were greatly reduced in the CNS of mice with EAE during CSF-1R inhibition. Given that monocyte recruitment into the CNS via CCL2/CCR2 signaling is essential to EAE pathology [53, 54], and that several reports have shown that CSF-1 induces CCL2 production by monocytes [55–57], we examined CCL2 production in the CNS of BLZ945- and anti-CSF-1-treated mice with EAE. The vast majority of CCL2^+^ cells were CD45^+^ (Supplemental Fig. 8A). Among CD45^+^ cells, there was a reduction in numbers of CCL2^+^ cells in both BLZ945- and anti-CSF-1-treated animals (Supplemental Fig. 7A, D). The majority of CCL2^+^ cells were CD11b^+^Ly6C^+^ (Supplemental Fig. 7B,C,E,F), indicating that monocyte-derived cells are a relevant source of CCL2 in the CNS during EAE. Most CCL2^+^ cells were TNF^+^MHC II^+^ inflammatory myeloid cells (Supplemental Fig. 8B,C,F,G). Notably, MFI for CCL2 among CCL2^+^ cells from anti-CSF-1-treated mice, but not from BLZ945-treated mice, was also reduced (Supplemental Fig. 8E). We also examined CCR2^+^ cells from the CNS of BLZ945- and anti-CSF-1-treated mice. As with CCL2-producing cells, there was a reduction in numbers of CCR2^+^ cells (Supplemental Fig. 7G,J). Most CCR2^+^ cells were CD45^Hi^CD11b^+^Ly6C^+^ cells (Supplemental Fig. 7H,I,K,L), indicating that these were the same cells that produce CCL2. Indeed, nearly all CCL2^+^ cells were CCR2^+^CD11b^+^ cells in anti-CSF-1-treated mice (Supplemental Fig. 8K). Combined with our *in vitro* findings, these data suggest that antagonism of CSF-1R signaling inhibits the survival and proliferation of monocytes/moDCs, resulting in fewer CCL2-producing cells, which then reduces the recruitment of CCR2^+^ cells in the CNS during EAE.

### Monocytes remaining in the CNS of anti-CSF-1-treated mice have a transcriptional profile consistent with a pro-survival phenotype

Anti-CSF-1 treatments depleted most (>80% depletion) but not all monocytes and monocyte-derived cells in the CNS of mice with EAE (Supplemental Fig. 10A,B). To identify transcriptional changes that could have enabled some monocytes to persist despite diminished CSF-1 signaling, we sequenced their transcriptome after 6 days of treatment, a timepoint that correlated well with maximal disease suppression (gating strategy shown in Supplemental Fig. 9A). There were 412 genes differentially expressed between monocytes from anti-CSF-1- and control MAb-treated mice (Supplemental Fig. 10C,D). We utilized the DAVID bioinformatics database [58, 59] to identify gene ontology, and KEGG pathway terms that were significantly enriched among the differentially expressed genes. Among GO terms identified as significantly enriched, the largest percentage of genes were involved in cell division (Supplemental Fig. 10E and Supplemental Fig. 9B). Among enriched KEGG pathways in these monocytes, the module with the greatest number of genes was the PI3K-Akt signaling pathway (Supplemental Fig. 10F and Supplemental Fig. 9C), which controls proliferation [60]. Among genes in this pathway, a number of growth factor receptors and transcription factors were upregulated, including *Flt1* (VEGFR), *Myb*, *Kit*, *Pdgfrb*, and *Fgfr1* (Supplemental Fig. 10G). These data suggest that monocytes in the CNS of anti-CSF-1-treated mice survive by upregulation of alternative growth factor receptors, which compensate for diminished CSF-1R signaling.

### BM-derived immune cells are major contributors to CSF-1R-dependent pathology in EAE

Inhibition of CSF-1R signaling during EAE, either by direct blocking of CSF-1R kinase activity or by neutralizing CSF-1, diminished numbers of moDCs and microglia, suggesting that disease suppression was due to reduced numbers of one or both of these cell types. To further test the role of CSF-1R signaling in monocytes and microglia in EAE, we developed a genetic model for *Csf1r* deletion that circumvents the perinatal lethality of conventional *Csf1r* knockout mice [61]. We crossed UBC-CreERT2 mice [62] and *Csf1r*^fl/fl^ mice [63] to generate CSF1RiKO mice which have tamoxifen-inducible Cre-mediated deletion of *Csf1r*. Tamoxifen-treated adult CSF1RiKO mice had 70%-100% reduced CSF-1R protein in western blots of cell lysates from the spleen, dLN and CNS (Supplemental Fig. 11A). Next, we treated CSF1RiKO mice with tamoxifen for 5 days and rested them for additional 7-14 days before immunization to induce EAE. CSF1R-iKO mice developed milder EAE than control mice, without a delay in disease onset (Fig. 6A,B, Supplemental Fig. 11B). The tamoxifen pre-treatment led to sustained reduction of CNS moDCs for the duration of observation (∼30 days p.i.) (Fig. 6C,D). In contrast, microglia were initially efficiently depleted, but by ∼30 days p.i. microglia numbers had recovered to near normal, indicating incomplete knockout of *Csf1r* in microglia progenitors. Nevertheless, microglia were substantially reduced at the time of immunization and until disease peak (∼day 20 p.i.). Thus, CSF1RiKO mice enable efficient depletion of CSF-1R in adult mice without the developmental defects present in conventional CSF-1R knockout mice; the depletion of moDCs is long lasting, whereas the depletion of microglia is transient.

**Figure 6.**
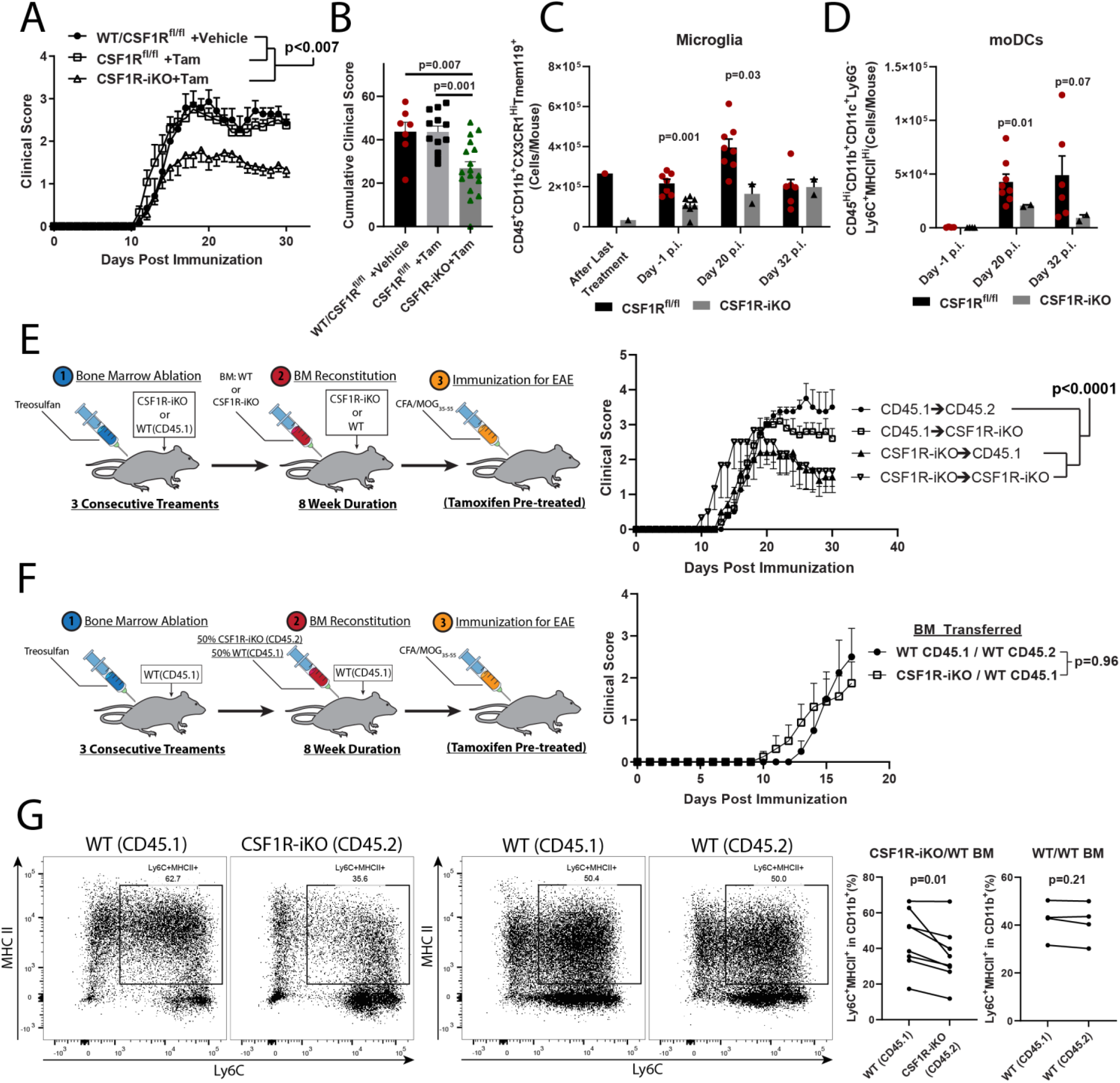
BM-derived immune cells are the major contributors to CSF-1R-dependent pathology. **A)** Comparison between vehicle (n=7) and tamoxifen-treated (n=17) *Csf1r*^fl/fl^ mice and tamoxifen-treated CSF1R-iKO mice (n=12). **B)** Quantification of cumulative score from C). Significance for clinical course determined by two-way repeated measures ANOVA, and by unpaired Student’s *t* test for cumulative scores. **C)** Time course showing numbers of microglia and **D)** moDCs in tamoxifen-treated *Csf1r*^fl/fl^ and CSF1R-iKO mice. Significance was calculated by unpaired Student’s *t* test **E)** BM was ablated with treosulfan, and 10^7^ WT or CSF1R-iKO BM cells were i.v. transferred into either WT or CSF1R-iKO recipients. After 8 weeks reconstitution, mice were treated with tamoxifen and immunized for EAE. Right panel shows clinical course of EAE. Numbers of mice per group were 4, 5, 5, 3 for group listed in legend from top to bottom. **F)** Mixed BM chimera experiments. BM was ablated as in G, then equal numbers of WT and CSF1R-iKO BM cells were co-transferred to recipients (total cell number transferred = 10^7^ cells/mouse) via i.v. injection. After reconstitution period, mice were treated with tamoxifen, rested, and then immunized for EAE induction. Significance for BM chimera experiments was calculated by two-way repeated measures ANOVA. **G)** Quantification of frequency of monocyte-derived APCs among CD11b^+^ cells from mixed BM chimeras in H). Significance was determined by two-tailed paired *t* test.

To determine the relative contributions of CSF-1R signaling in monocytes and microglia to EAE, we generated BM-chimeras by reconstituting treosulfan-conditioned WT and CSF1RiKO recipient mice with either WT or CSF1RiKO BM. We used the BM-conditioning agent treosulfan because it does not cross or affect the BBB, does not induce a cytokine storm, nor does it enable engraftment of peripheral myeloid cells into the CNS, as has been observed in irradiation-induced BM chimeras [64–66]. Pre-conditioning with treosulfan was highly efficient for establishing chimerism, with >90% of peripheral immune cells in chimera mice being of donor BM origin (Supplemental Fig. 11C). Following 8 weeks of BM-reconstitution, mice were treated with tamoxifen to knockout *Csf1r* and then immunized to induce EAE (Fig. 6E). Both WT and CSF1R-iKO mice reconstituted with WT BM developed typical EAE, whereas WT mice reconstituted with CSF1RiKO BM developed attenuated EAE, similar to CSF1RiKO mice reconstituted with CSF1RiKO BM. These data show that BM-derived cells are a major contributor to CSF-1R-dependent pathology during EAE, whereas microglia play a less important role.

Lastly, we sought to understand the role of CSF-1R in survival of monocytes during EAE. We utilized mixed-BM chimeras generated by reconstitution of WT mice with WT and CSF1R-iKO BMs (1:1). 50% WT BM is sufficient to drive the development of typical EAE [6, 67]. Indeed, mice reconstituted with a mixture of WT and CSF1R-iKO BMs and pre-treated with tamoxifen developed EAE equivalent to mice reconstituted with WT BM (Fig. 6F). In the CNS, we found a lower frequency of CSF1R-iKO monocyte-derived cells when compared to WT cells (Fig. 6G, Supplemental Fig. 11D). These data suggest that CSF-1R signaling maintains monocytic cells in the CNS during EAE.

## DISCUSSION

We show that blocking CSF-1R, CSF-1 and IL-34 has differential effects on EAE. Overall, blocking either CSF- 1R or CSF-1 resulted in suppression of clinical disease and diminished CNS inflammation and demyelination. Notably, blocking CSF-1R and CSF-1 produced distinct effects on the composition of immune cells in the CNS during EAE. Numbers of microglia and monocyte-derived cells were the most reduced by CSF-1R inhibition. Treatment with BLZ945 depleted virtually all microglia, whereas anti-CSF-1 treatment preferentially depleted inflammatory microglia, reducing their number to similar numbers as in naïve mice. Moreover, depletion of myeloid cells in anti-CSF-1-treated mice was limited to CNS lesions and adjacent areas, while grey matter microglia had a similar appearance as naïve mice. The notably lower microglia depletion by anti-CSF-1 MAb than with BLZ945 did not result in less effective disease suppression, but rather improved long-term therapeutic efficacy when compared to BLZ945 treatments, suggesting that full therapeutic benefit can be achieved without applying a maximally ablative approach. Limited depletion of microglia by anti-CSF-1 MAb may therefore be advantageous in MS therapy, as it carries fewer potential risks than widespread microglia depletion likely would. Favoring a milder approach for depletion of myeloid cells is supported by a finding that PLX5622, a commonly used inhibitor of CSF-1R kinase activity, in addition to depleting microglia, also causes long-term and widespread systemic changes to myeloid and lymphoid compartments [68]. It is likely that other small molecule inhibitors of CSF-1R, including BLZ945, induce similar changes, but it is also likely that blocking CSF-1 induces fewer systemic changes because of compensatory IL-34 signaling and more restricted tissue distribution of the MAb compared to small molecule inhibitors, as exemplified by our findings in the CNS. For example, other organs with blood barriers, such as testes, eye, and thymus, all of which express CSF-1R [69], would likely be less affected by anti-CSF-1 MAb than a small molecule inhibitor, which could more readily penetrate the barriers and inhibit both CSF-1 and IL-34 signaling.

The limited microglia depletion by anti-CSF-1 MAb is likely due to the presence of IL-34 in non-inflamed CNS areas, where it maintains homeostatic microglia survival. Indeed, it has been shown that systemic injections of anti-CSF-1 MAb do not deplete microglia [70]. This is consistent with studies showing that IL-34 accounts for approximately 70% of CSF-1R signaling in healthy brain [23], and that anti-CSF-1 MAb is unlikely to penetrate into the normal CNS parenchyma, as only a miniscule fraction of Abs crosses the intact BBB [10, 30]. Thus, it is expected that neutralization of CSF-1 in the CNS occurs primarily in active lesions, where BBB is leaky, resulting in localized depletion of inflammatory myeloid cells. This model is analogous to the role of CSF-1R ligands in the skin, where IL-34 maintains Langerhans cells in steady state. However, during skin inflammation, IL-34 becomes dispensable, as infiltrated immune cells produce CSF-1 and maintain/expand numbers of Langerhans cells [45]. It should be noted, though, that CSF-1R signaling is not in itself inherently pro-inflammatory by eliciting inflammatory phenotype in myeloid cells, but can have such a net effect by simply maintaining their survival during inflammation. In fact, in the absence of inflammation, CSF-1R signaling induces an immunosuppressive/homeostatic M2 phenotype in macrophages and a resting/quiescent phenotype in microglia [1, 12–26]. Hence, our observations on blockade of CSF-1R signaling in EAE are the net effect of abrogating both pro- and anti-inflammatory functions of CSF-1R signaling, with the pro-inflammatory ones predominating. Together, our findings suggest that CSF-1 promotes inflammation in EAE by expansion of microglia and monocyte-derived myeloid cells, whereas IL-34 maintains microglia in non-inflamed CNS areas, similar to the healthy CNS.

Our data suggest that IL-34 does not play a significant role in EAE pathology. This can be explained by notably more widespread and abundant expression of CSF-1 compared with IL-34 [71]. It is likely that in most cases CSF-1 can therefore compensate for lack of IL-34. This interpretation is supported by a study that has demonstrated that CSF-1 in inflamed sites becomes the dominant CSF-1R ligand, even in tissue (skin) where IL-34, but not CSF-1, is expressed in steady state [45]. Thus, although our data suggest that blockade of IL-34 with MAb may have been incomplete due to the development of anti-rat IgG responses, it is probable that abundantly produced CSF-1 in CNS lesions mediates most CSF-1R signaling, and that blockade of IL-34 therefore does not have an effect on EAE.

Our in vitro studies with BMDCs show that the primary effect of CSF-1R inhibition is limiting the number of myeloid DCs in these cultures rather than affecting their APC functions. These data are consistent with a body of literature showing that CSF-1R signaling in myeloid cells chiefly provides proliferative and anti-apoptotic signals for maintenance of the population size [34]. This concept is exemplified by differences between animals lacking CSF-1 and IL-34, in which CSF-1 knockout mice have reduced numbers of osteoclasts and monocytes but only a modest reduction in microglia [32]. In contrast, IL-34 knockout mice have greatly reduced numbers of microglia and Langerhans cells, but largely normal numbers of other tissue resident macrophages [32]. Thus, CSF-1R inhibition is likely to suppress inflammation in EAE by reducing the population size of inflammatory myeloid cells in the CNS.

Transcriptional profiling of monocytes that remained in the CNS after anti-CSF-1 treatment suggests that they avoid death via upregulation of growth factor receptors known to promote myeloid cell survival, including Kit [72–74]. Notably, upregulation of these genes has been reported in myeloid cell cancers [75–79]. However, a number of questions regarding these surviving monocytes remain, such as: are they a normally present subpopulation among CNS monocytes, or were they induced by neutralization of CSF-1; do they eventually succumb to early death compared to monocytes that have been receiving CSF-1R signaling; do all monocytes lacking CSF-1R signaling acquire this phenotype before death; and, what is the capacity of the surviving monocytes to perpetuate inflammation? It is possible that altered phenotype of surviving monocytes is less pro-inflammatory because of diminished effector functions, such as cytokine and chemokine production.

Our in vivo studies indicate an additional mechanism of EAE suppression by inhibition of CSF-1R signaling, namely, reduced recruitment of immune cells to the CNS. We found reduction in numbers of both CCL2- and CCR2-expressing cells when mice were treated with BLZ945 or anti-CSF-1 MAb. Most cells that expressed CCL2 and CCR2 were monocytes/moDCs, which suggests a model whereby monocytes that infiltrate the CNS produce CCL2, thus amplifying inflammation by further recruitment of CCR2-expressing cells. Given that CSF-1R signaling in monocytes/macrophages induces CCL2 expression [55–57], this suggests that its blockade reduces CNS inflammation by two mechanisms: 1) by reducing numbers of cells that produce CCL2, and 2) by reducing CCL2 production of surviving cells, which together amounts to greatly diminished CCL2 levels in the CNS during EAE. This is consistent with an essential role of CCL2 and CCR2 in EAE, since interventions that affect them attenuate disease [80–82]. It is also possible that in addition to monocytes, reduction in CCL2 directly affects recruitment of pathogenic CCR2^+^ Th cells to the CNS, given a report that CCR2 drives their recruitment to the CNS [80]. Taken together with our result showing reduced GM-CSF and IL-1β production in the CNS, it is likely that CSF-1R inhibition suppresses EAE by depleting CCL2- and IL-1β-expressing APCs. IL-1β has an essential role in EAE [6, 48, 83, 84] by acting on CD4^+^ T cells to promote their proliferation and GM-CSF production; GM-CSF is also essential to EAE development by acting on monocytes to induce their pro-inflammatory phenotype and IL-1β production, thus completing a positive feedback loop that sustains inflammation [85]. Inhibition of CSF-1R signaling likely interrupts this pro-inflammatory feedback loop, resulting in EAE suppression.

To determine the relative contributions of CSF-1R signaling in monocytes and microglia to EAE, we developed CSF1R-iKO mice. Our BM chimera experiments indicate that a lack of CSF-1R signaling in BM-derived cells, rather than in CNS-resident cells (e.g. microglia), is responsible for the EAE suppression. However, the transient nature of microglia depletion in CSF-1RiKO mice prevents us from concluding that microglia are entirely dispensable for EAE development. Our experiments with mixed BM chimeras support the view that CSF-1R functions as a growth factor receptor for monocytes and their progeny [9–11]. However, it is worth noting that treating WT mice with BLZ945 induced more profound reduction in CNS moDCs than in mixed BM chimeras. This may be explained by the presence of WT monocytes in mixed BM-chimeras, which are able to produce CCL2, and therefore maintain recruitment of monocytes into the CNS. This is consistent with a model in which CSF-1R signaling maintains numbers of monocytes and cells that differentiate from them, by promoting their survival/proliferation, but also by potentiating their recruitment into the CNS via CCL2 production, which are both reported functions of CSF-1R [9–11, 55–57].

Therapies using anti-CSF-1 MAb and small molecule inhibitors of CSF-1R have been tested in multiple clinical trials for autoimmune and oncological diseases [86–91]. These trials have demonstrated that blockade of CSF-1/CSF-1R is well tolerated by patients [90, 92, 93]. Targeting CSF-1/CSF-1R in MS has not been tested, but agents used in trials for other diseases would likely be suitable for testing in MS. Moreover, because CSF-1R signaling is not required by myeloid progenitors residing in the BM or CNS (for microglia) [34, 94], the effects of these treatments would be largely reversible. Indeed, there is complete repopulation of microglia within one week after cessation of treatment with CSF-1R inhibitors [94]. A treatment modality can be envisioned whereby blocking CSF-1R signaling for therapy of MS would follow an intermittent regimen, given for a period of time, instead of continuously. Thus, as a potential therapy for MS, targeting CSF-1/CSF-1R offers several advantages, including the potential of being readily translatable to clinical testing with already existing therapeutic agents.

In conclusion, blocking CSF-1R signaling ameliorates EAE by depleting inflammatory APCs in the CNS. MS therapy with anti-CSF-1 MAb could be a preferred approach, because, unlike small molecule inhibitors of CSF-1R, it preserves quiescent microglia and their homeostatic functions, as well as other IL-34 functions, such as maintenance of Langerhans cells. Limited depletion of microglia by anti-CSF-1 treatment, however, does not diminish its therapeutic effect compared to BLZ945 treatment. Reducing CSF-1R signaling via neutralization of CSF-1 could therefore be a novel strategy for therapy of MS.

## MATERIALS AND METHODS

### More information is available in supplemental materials

#### Mice

All mice used in this study were on C57BL/6J genetic background. Mice were either obtained from The Jackson Laboratories (Bar Harbor, Maine) or bred in-house. All experimental procedures were approved by the Institutional Animal Care and Use Committee of Thomas Jefferson University. CSF1RiKO mice were generated by breeding UBC-CreERT2 (The Jackson Laboratories, Stock #007001) with Csf1r^flox^ mice (The Jackson Laboratories, Stock #021212). Genotyping was performed as recommended by The Jackson Laboratories. CD45.1 mice (B6.SJL-Ptprca Pepcb/BoyJ, Stock #002014) were obtained from The Jackson Laboratories.

#### Induction and Scoring of EAE

EAE was induced by immunization with 1:1 emulsion of PBS and complete Freund’s adjuvant (CFA) containing 5 mg/mL heat killed *M. tuberculosis* (BD Biosciences) and 1 mg/mL MOG_35-55_ peptide (Genscript). Mice were immunized on both flanks by subcutaneous injection of the emulsion for a total of 200 µL. Pertussis toxin was i.p. injected on days 0 and 2 post-immunization at 200 ng per dose. Mice were scored according to the following scale: 0 - No clinical symptoms. 0.5 - Partial paralysis of the tail or waddling gait. 1.0 - Full paralysis of the tail. 1.5 - Full paralysis of the tail and waddling gait. 2.0 - Partial paralysis in one leg. 2.5 - Partial paralysis in both legs or one leg paralyzed. 3.0 - Both legs paralyzed. 3.5 - Ascending paralysis. 4.0 - Paralysis above the hips. 4.5 – Moribund; mouse being unable to right itself for 30 seconds. 5.0 - Death.

#### BLZ945 Preparation and Treatment

BLZ945 (Selleck Chemicals and MedChemExpress) was prepared from powder in 20% captisol at 12 mg/mL. Mice were treated with BLZ945 by oral gavage with 4-6 mg/treatment/day. In initial experiments, we used BLZ945 prepared in 20% captisol, and 20% captisol as control, which were a generous gift from Novartis International AG.

#### *In vivo* CSF-1, Anti-CSF-1, Anti-IL-34, and Anti-CSF-1R Treatments

Recombinant CSF-1 (4 µg/dose; R&D Systems) was given to EAE mice on days 4, 8, 12, 16 post immunization (p.i.) by intraperitoneal (i.p.) injection. All MAb treatments were also given by i.p. injection. Prophylactic treatments with anti-CSF-1 (200 µg/dose; clone: 5A1; Bio X Cell) started on day 0 p.i. and were given every other day until disease onset, when dosing was changed to every day. In therapeutic treatments, MAb was given every day, starting on days indicated in figures, for the duration of acute phase of the disease (typically days 11-25 p.i.), then switched to every other day for the rest of the experiments. Equal amounts of control IgG1 (Clone: HPRN; Bio X Cell) were used to treat control mice. Mice were treated with anti-IL-34 MAb (100 µg/dose; Clone: 780310; Novus Biologicals) every other day for the duration of the experiment. Anti-CSF-1R Mab (400 µg/dose; Clone: AFS98; Bio X Cell) was given every other day. In experiments with anti-IL-34 and anti-CSF-1R MAbs, equal amounts of control IgG2A (Clone: 2A3; Bio X Cell) were given to control mice.

## Author Contributions

Bogoljub Ciric and Daniel Hwang were responsible for the conceptualization of this project. Daniel Hwang was responsible for carrying out experiments and analysis of data.

Larissa Lumi Watanabe Ishikawa, Alexandra Boehm, Ziver Sahin, Giacomo Casella, Soohwa Jang, and Maryamsadat Seyedsadr assisted in carrying out experiments.

Michael Gonzalez, James Garifallo and Hakon Hakonarson were responsible for RNA sequencing experiments and assisted with analysis of RNA sequencing data.

Guang-Xian Zhang, Weifeng Zhang, Dan Xiao, and Abdolmohamad Rostami assisted with manuscript preparation and provided helpful insights in interpreting data.

## Acknowledgments

We thank Katherine Regan for editing the manuscript. We thank Medac Germany for the kind gift of treosulfan for our BM chimera studies.

## SUPPLEMENTAL MATERIALS

### MATERIALS AND METHODS

#### Flow Cytometry

Isolated cells were stimulated with PMA (500 ng/mL; Sigma Aldrich), ionomycin (50 ng/mL; Sigma Aldrich), and 1 µL/mL Golgiplug (BD Biosciences) for 4 h at 37°C. After stimulation, cells were washed with PBS containing 3% FBS (v/v). Cell surface antigens were stained with Abs in 100 µL of PBS/3% FBS for 20-30 min at 4°C. Cells were then washed and fixed with 100 µL Fix and Perm Medium A (Thermo Fisher) for 20 min at room temperature and washed again. Cells were permeabilized with Fix and Perm Medium B (Thermo Fisher) and stained with Abs against intracellular antigens in 100 µL Fix and Perm Medium B and 100 µL PBS/3%FBS for 1 h. Cells were then washed twice, resuspended in 500 µL PBS and analyzed on a BD FACSAria Fusion flow cytometer (BD Biosciences).

#### Isolation of Immune Cells from the CNS

Mice were anesthetized and blood was removed by perfusion with 60 mL PBS. Spinal cord was flushed out of the spinal column with PBS. Brains and spinal cords were pooled and cut manually into small pieces in 700 µL Liberase TL dissolved in RPMI at 0.7 mg/mL (Roche) then incubated at 37°C for 30 min before reaction was quenched using complete media containing FBS. Tissue was homogenized by pushing through a 100 µm sterile filter with syringe plunger. Homogenate was centrifuged at 1500 RPM (300 x g) for 5 min and resuspended in 25 mL of 70% 1x Percoll-PBS (90% Percoll, 10% 10x PBS). 25 mL of 30% Percoll-PBS was gently overlayed onto the 70% layer and was centrifuged at 2000 RPM without brake at room temperature for 30 min. Cells that pooled at the interface of 30/70% layers and the majority of the 30% layer were then collected, diluted with PBS or media and centrifuged at 1500 RPM (300 x g) for 5 min.

#### Antibody Titer Measurement

Serum was collected from peripheral blood of animals treated with Rat IgG_2A_ (Clone: 2A3; Bio X Cell), anti-CSF1 MAb (Clone: 5A1; Bio X Cell), anti-CSF-1R MAb (Clone: AFS98; Bio X Cell), anti-IL-34 MAb (Clone: 780310, Novus Biologicals). 96-well ELISA plates were coated with the same MAb that was used to treat animals overnight at room temperature and blocked with 1% BSA for 2 h. Plates were washed and incubated with serum for 1 h and washed. Anti-Mouse IgG-HRP conjugate secondary Ab (Jackson Immunolabs) was used to detect the presence of anti-Rat IgG response by measuring absorbance at 450 nm and subtracting absorbance at 540 nm.

#### Library Preparation and RNA-seq Analyses

Next-generation sequencing libraries were prepared using the Illumina TruSeq Stranded Total RNA library preparation kit, with high quality RNA (RIN >= 8.7) and 200 ng of input RNA. Libraries were assessed for quality using the PerkinElmer Labchip GX and qPCR using the Kapa Library Quantification Kit and the Life Technologies Viia7 Real-time PCR instrument. Libraries were diluted to 2 nM and sequenced in a paired-end (2 x 100bp), dual-indexed format on the Illumina HiSeq2500 using the High Output v4 chemistry.

RNA-seq reads were demultiplexed into sample-specific fastq files and aligned to the mm10 reference genome using the DRAGEN genome pipeline [95] to produce BAM files. Generated BAM files were read into R statistical computing environment and gene counts were obtained using the Rsubread package, producing a feature/gene counts matrix. Differential expression analysis was performed using the R/Bioconductor package DESeq2, which uses a negative binomial model [96]. Analysis was performed using standard thresholds and parameters while filtering genes with low mean normalized counts. Further downstream analysis was performed using the Ingenuity Pathway Analysis (IPA) and GSEA software using the normalized read count table. Additionally, differentially expressed genes (p<0.01) were entered into the DAVID utility for functional annotation and analyzed for gene ontology terms for biological processes and for KEGG pathway terms [58, 59].

#### Tamoxifen Treatment

Tamoxifen (Sigma) was resuspended in corn oil (Sigma) to 20 mg/mL by heating in a 37°C water bath and vortexing. For tamoxifen treatment via i.p. injection, 2 mg of tamoxifen was injected per day for a total of 5 injections. Mice were then rested for 14-21 days before further manipulation. For tamoxifen treatment via oral gavage, mice were treated 5 times with 5 mg of tamoxifen per day, then rested for 7 days.

#### Immunohistochemistry and Immunofluorescence Microscopy

Animals were perfused with 50 mL PBS followed by 40 mL cold paraformaldehyde (PFA) solution. The spinal column was then removed and placed in PFA overnight. The day after perfusion, tissue was transferred to 30% sucrose in PBS and allowed to incubate overnight, or until tissue had sunk to the bottom of the tube. Spinal cord was then dissected from the spinal column, cut into 4 pieces of approximately equal length and embedded in OCT. The entire spinal cord was sectioned into 10 µm-thick coronal sections, skipping 100 µm between each section. Prior to staining, sections were immersed in PBS for 10 min to remove residual OCT. For Sudan black staining, sections were stained by immersion for 30 min in a 10 mg/mL solution of Sudan black prepared in 70% ethanol. Following staining, slides were decolorized by immersion in a 0.1% Triton-X100 in PBS solution for 30 min. For immunofluorescence experiments, slides were blocked using 10% Normal Goat Serum with 0.1% sodium azide (Thermo Fisher) for 30 min. Primary Abs for Iba1 (1:500 dilution, SySy, Cat# 234 006), and CD68 (1:100 dilution, Thermo Fisher, Clone: FA-11) were diluted in 10% Normal Goat Serum with 0.3% Triton-X100. Slides were stained with primary Abs overnight at 4°C, then washed 3 times with PBS for 5 min. Secondary Abs (Thermo Fisher) were diluted 1:1000 and slides were stained for 1 h. Slides were then washed 3 times with PBS, mounted with ProLong™ Diamond Antifade Mountant with DAPI (Thermo Fisher) and imaged using a Nikon AR1 confocal microscope at the Thomas Jefferson University Imaging core facility.

#### Analysis of Immunofluorescence Microscopy

Demyelination in Sudan black-stained spinal cord sections was determined by dividing total demyelinated area of white matter by total white matter area, using NIS elements software (Nikon). Immunofluorescence images were analyzed using Slidebook 6 software (Intelligent Imaging Innovations). Quantification of cell density was performed by masking cell nuclei by fluorescent thresholding. Touching nuclei were separated by applying the watershed algorithm at 25% aggressiveness and then again at 50% after removal of small objects. Errors in object identification or over-segmentation caused by the watershed algorithm were then corrected manually. Features of spinal cord sections were manually masked (e.g. lesions, grey matter, etc.) and cells not belonging to that feature were discarded by only keeping cells that overlapped with the feature mask. Object IDs were then assigned to individual cells and mean pixel intensity per object was determined for each channel. A threshold intensity value was then assigned for each channel and used to determine positivity for that marker. The number of positive cells was then divided by the area of that feature to determine cell density.

#### Western Blot

Cells were re-suspended in a RIPA lysis buffer supplemented with a protease inhibitor cocktail (Sigma-Aldrich). 30 mg proteins diluted with Laemmli buffer and loaded onto 8%–14% polyacrylamide gels. Protein concentrations were measured with bicinchoninic acid (BCA) assay (Pierce). Anti-CSF-1R (Abcam) and mouse anti-β actin (Sigma) were used as primary antibodies.

#### Bone Marrow-Derived DC Culture

Bone marrow (BM) was isolated from tibia and femurs of mice, and BM cells were cultured at 1 x 10^6^ cells per mL in a total volume of 10 mL in petri dishes with GM-CSF (20 ng/mL) + IL-4 (20 ng/mL) for 4 days. On the 4^th^ day, 5 mL of media was removed from the plates, cells were pelleted by centrifugation and resuspended in 5 mL fresh media containing GM-CSF and IL-4. The cell suspension was added back to the original petri dishes and cultured for an additional 3 days. To induce a mature DC phenotype, cells were washed, replated in fresh media containing LPS (300 ng/mL) and cultured for 72 h. For cultures involving monocytes, CD11b^+^ cells were isolated from BM cell suspension of CD45.1^+^ mice by MACS positive selection (Miltenyi Biotec), after which Ly6C^Hi^Ly6G^-^ monocytes were sorted on a FACSAria Fusion instrument (BD Biosciences). Isolated monocytes were then added to CD45.2^+^ total BM cell cultures.

#### Co-culture of DCs and T cells and Proliferation Assay

For proliferation assays, 4 x 10^4^ DCs were co-cultured with 1.6 x 10^5^ 2D2 CD4^+^ T cells in 96-well U-bottom tissue culture plates. Cells were stimulated with MOG_35-55_ peptide (25 μg/mL) for 72 h. At approximately 60 h after starting the culture, 1 µCi of [^3^H]Thymidine (Perkin-Elmer) was added to each well in a volume of 50 µL. Cells were harvested 24 h later and counts per minute (C.P.M.) measured in MicroBeta2 beta counter (Perkin-Elmer).

#### Bone Marrow Chimeras

Treosulfan (L-threitol-1,4-dimethanesulfonate; a generous gift from Medac Germany) was resuspended in sterile water at a concentration of 50 mg/mL and administered by i.p. injection over 3 consecutive days at 2000 mg/kg per dose. Mice were then rested for 72-96 h before i.v. transfer of 10^7^ BM cells.

### SUPPLEMENTAL FIGURES

**Supplemental Figure 1:**
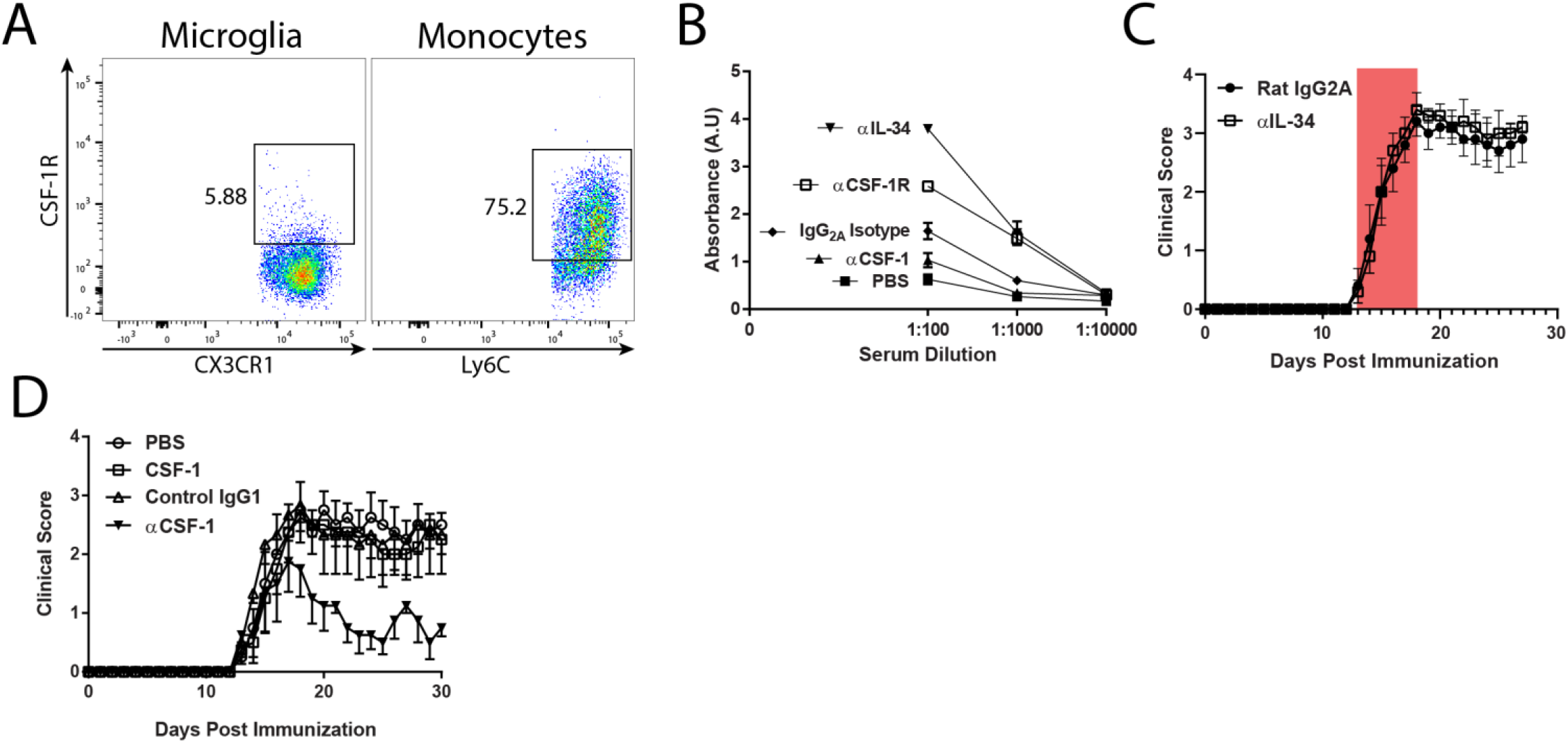
**A)** C57BL6/J mice were immunized with MOG35-55 and treated with anti-CSF1 MAb (200 µg, every other day), control rat IgG1 MAb (200 µg, every other day), recombinant mouse CSF-1 (4 µg per dose, given on days 4, 8, 12, 16 p.i.) or PBS by i.p. injection. **B)** Serum Ab titers of mice with EAE treated with MAbs against IL-34 (n=3), CSF-1R (n=3), IgG2a isotype (n=4), CSF1 (n=3), and control animals treated with PBS (n=6). Error bars are standard error from the mean. **C)** Flow cytometry plots depicting staining of CD45^+^CD11b^+^CX3CR1^Hi^ microglia and CD45^Hi^CD11b^+^Ly6G^-^Ly6C^Hi^ monocytes for CSF-1R with the AFS98 MAb. **D)** C57BL6/J mice were immunized with MOG35-55 and treated with anti-IL-34 MAb (55 µg per day, i.p) for the period indicated by the red box.

**Supplemental Figure 2:**
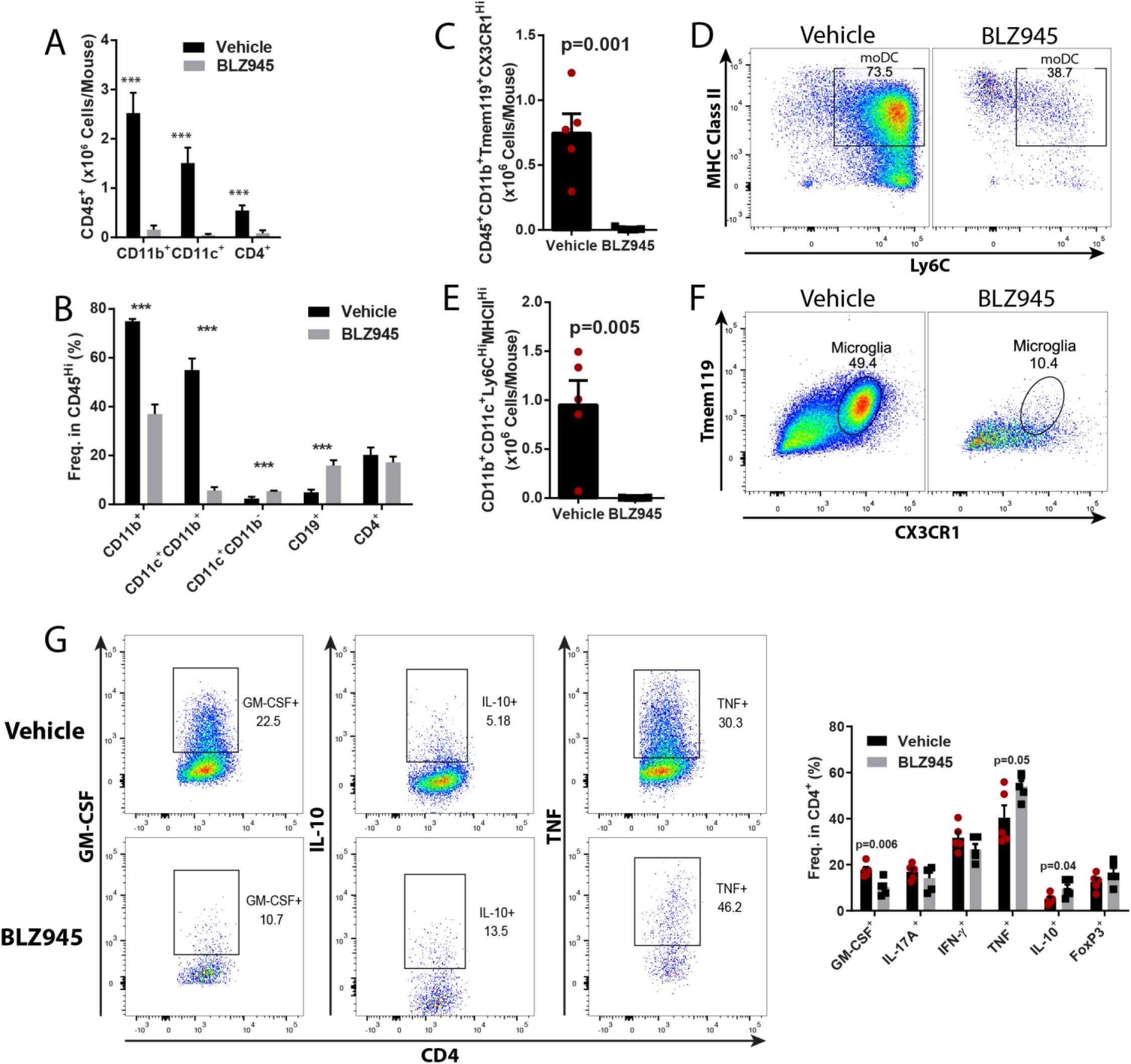
Inhibition of CSF-1R signaling alters both myeloid and T cell compartments in the CNS of EAE mice. C57BL/6J mice were immunized with MOG35-55 for EAE induction and treated orally with BLZ945 (200 mg/kg/day) or vehicle control (20% captisol) daily, starting on the day of immunization. Mice were sacrificed on day 15 p.i., and pooled brain and spinal cords of each mouse were used for cell isolation (n=5/group). **A)** Numbers of CD11b^+^, CD11c^+^ and CD4^+^ cells. **B)** Frequency of CD11b^+^, CD11b^+^CD11c^+^ CD11b^+^CD11c^-^, CD19^+^, and CD4^+^ cells in CD45^Hi^ cells. **C-F)** Numbers of CD45^+^CD11b^+^Tmem119^+^CX3CR1^Hi^ microglia and CD45^Hi^CD11b^+^Ly6C^Hi^MHCII^Hi^ moDCs. **G)** Quantification of GM-CSF, IL-17A, IFN-γ, TNF, IL-10 and FoxP3 expression by CD4^+^ cells from the CNS. Significance was calculated with Student’s *t* test. Error bars are S.E.M.

**Supplemental Figure 3.**
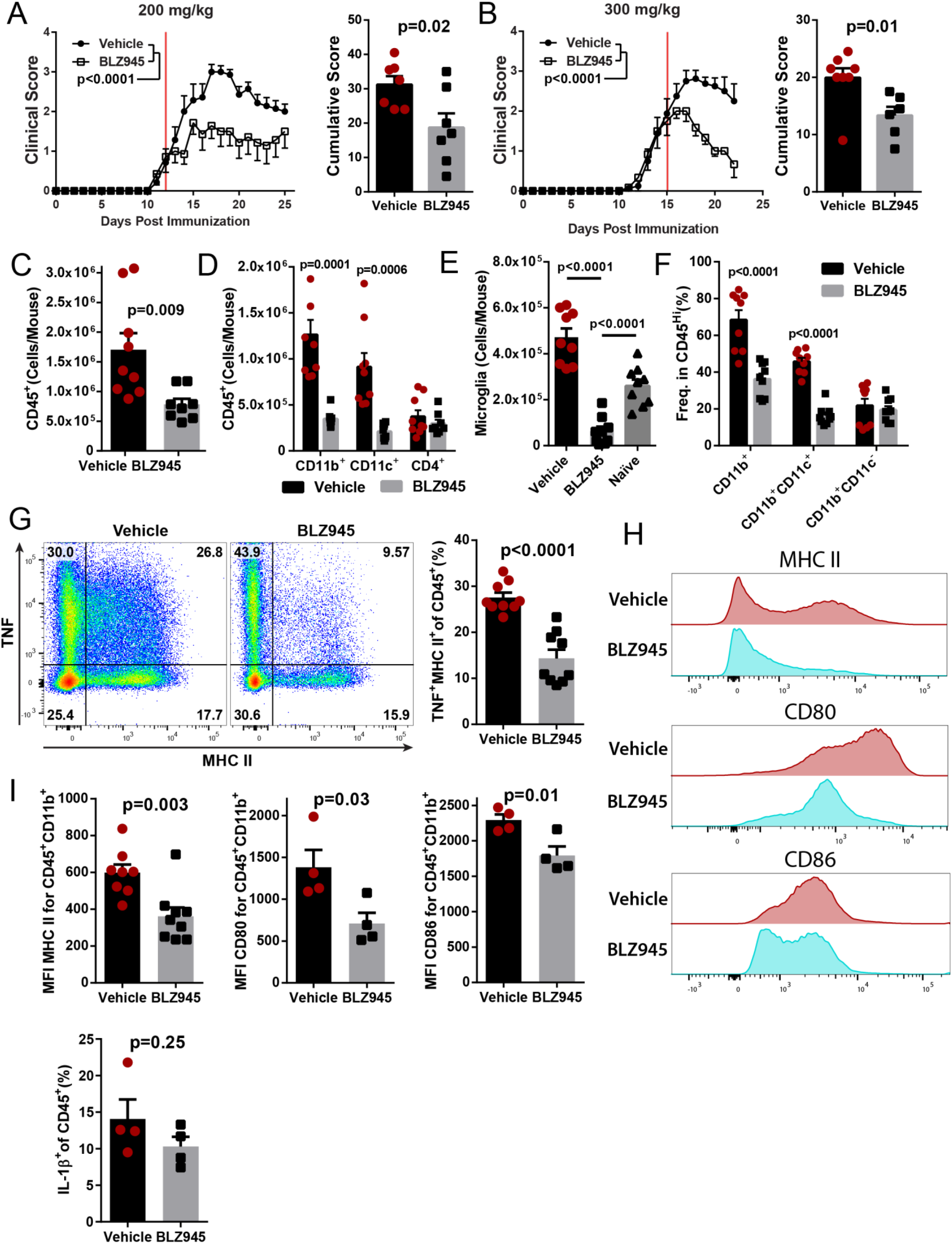
BLZ945 suppresses ongoing clinical EAE, and reduces the number of myeloid APCs in the CNS. C57BL/6J mice were immunized and allowed to develop clinical signs of EAE before treatment with BLZ945 (n=7) or vehicle (n=8). Treatments were given once per day by oral gavage. **A)** Clinical course and cumulative score for mice treated with 200 mg/kg BLZ945 starting at a clinical score of ∼1. Red line indicates start of treatment. **B)** Mice treated with 300 mg/kg BLZ945 (n=6) or vehicle (n=8), starting at a clinical score of ∼2. A) and B) were compiled from two independent experiments. Significance for clinical course determined by two-way repeated measures ANOVA, and by unpaired Student’s *t* test for cumulative scores. **C-I)** Analysis of the CNS (pooled brain and spinal cords) by flow cytometry. **C)** Number of CD45^+^ cells. **D)** Number of CD45^+^ cells that also expressed CD11b, CD11c, or CD4. **E)** Number of CD45^Lo^CD11b^+^CX3CR1^Hi^ microglia. Naïve mice are untreated WT C57BL/6J mice that were not immunized or otherwise manipulated. **F)** Frequency of CD45^Hi^ cells that also express CD11b and/or CD11c. **G)** TNF and MHC II expression in CD45^+^ cells. **H-I)** Expression of MHC II, CD80 and CD86 in CD45^+^CD11b^+^ cells. Significance for C-I) was calculated by unpaired Student’s *t* test. P-value corrections for multiple comparisons was performed by false discovery rate approach with Q=0.01 as a cutoff. Error bars are S.E.M.

**Supplemental Figure 4:**
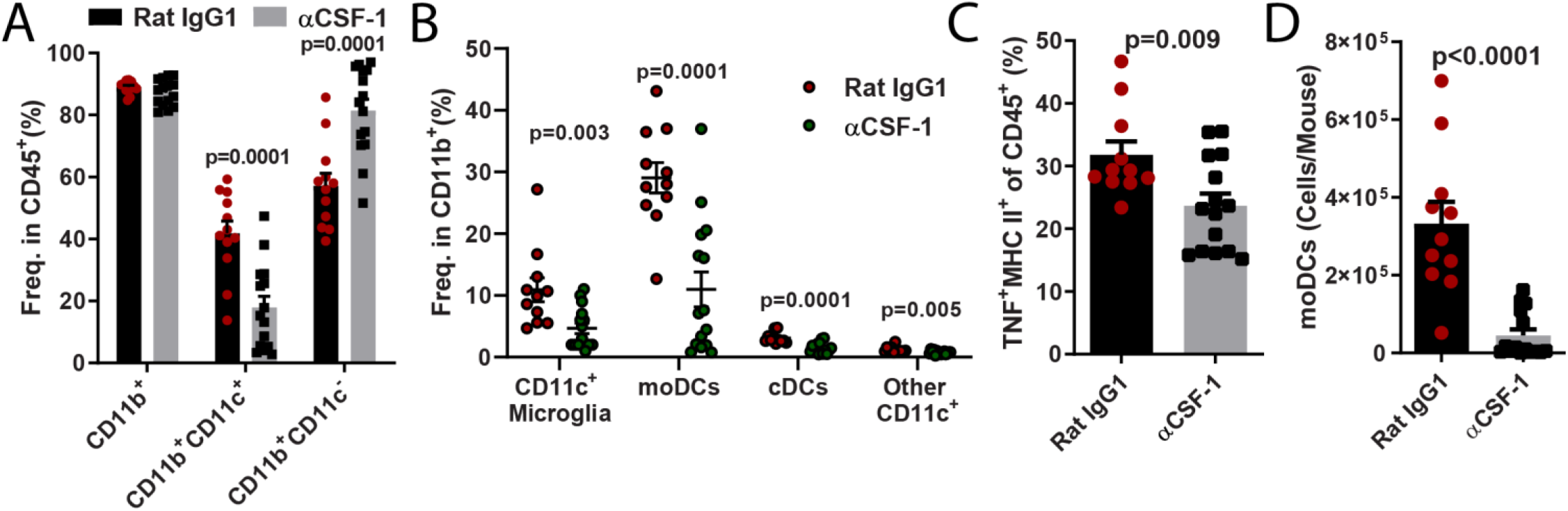
Anti-CSF-1 treatment reduces numbers of DCs in the CNS of mice with EAE. **A)** Frequency of CD11b^+^, CD11b^+^CD11c^+^, CD11b^+^CD11c^-^ cells among CD45^+^ cells from the CNS of mice with EAE treated with either anti-CSF-1 or control MAb. **B)** Frequency of CD11c^+^ microglia (CD45^+^CD11b^+^CX3CR1^Hi^Tmem119^+^), moDCs (CD45^Hi^CD11b^+^CD11c^+^Ly6G^-^Ly6C^Hi^MHCII^Hi^), cDCs (CD45^Hi^CD11b^+^CD11c^+^CD26^+^Ly6C^-^MHCII^+^) and other CD11c^+^ cells among CD11b^+^ cells. **C)** Number of CD45^Hi^CD11b^+^CD11c^+^Ly6G^-^Ly6C^Hi^MHCII^Hi^ moDCs. **D)** Frequency of TNF^+^MHC II^+^ cells among CD45^+^ cells. Significance was calculated with unpaired Student’s *t* test. Error bars are S.E.M.

**Supplementary Figure 5:**
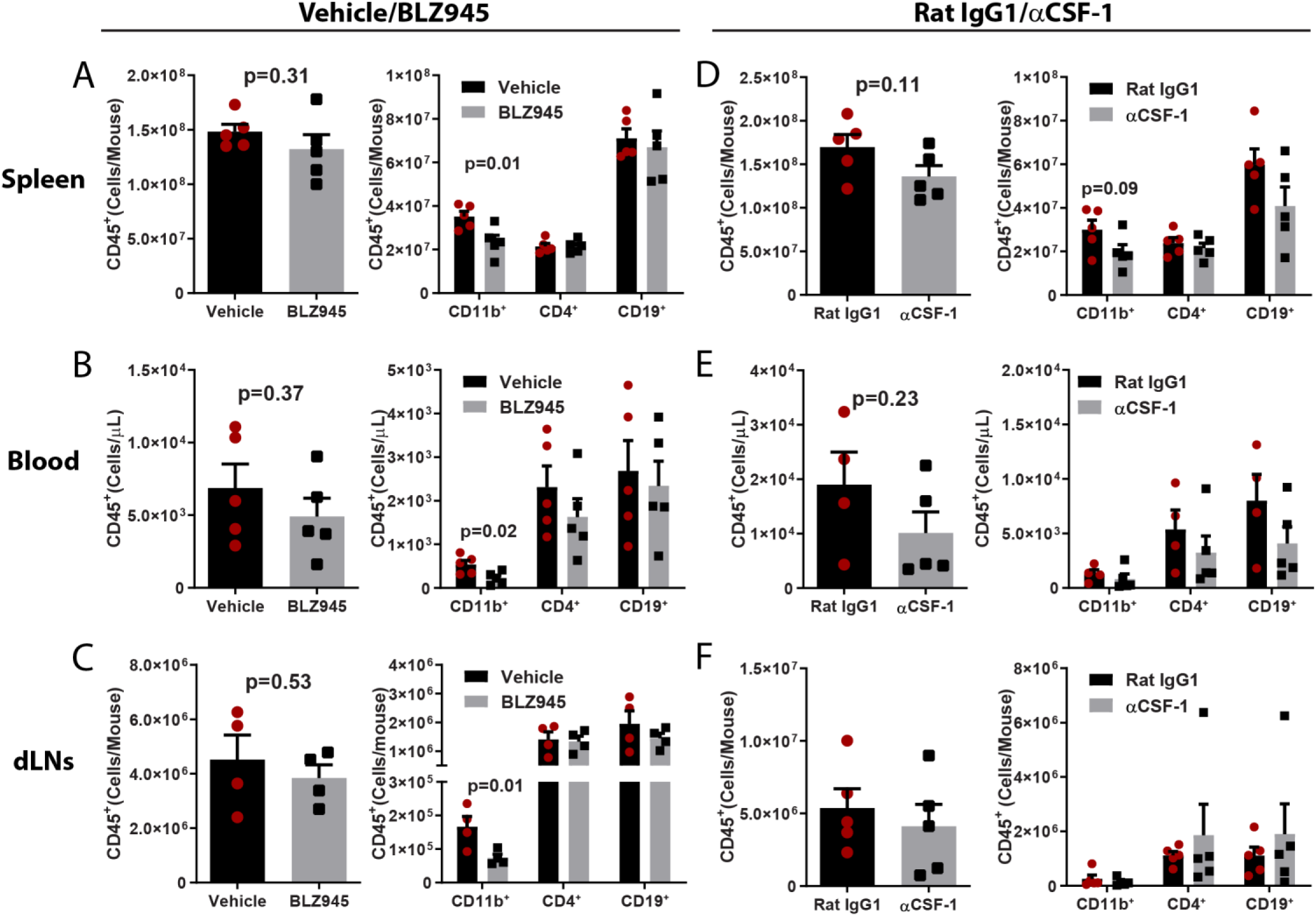
Immune response in peripheral lymphoid organs of BLZ945- and anti-CSF-1-treated mice on day 8 p.i. **A-C)** BLZ945- and vehicle-treated mice (n=5 per group). Number of CD45^+^, CD11b^+^, CD4^+^ and CD19^+^ cells in **A)** spleen, **B)** blood, and **C)** dLN. **E-F)** Same quantification as in A-C), but for anti-CSF-1- and rat IgG1 control MAb-treated mice. Statistical significance was calculated using two-way unpaired *t* test. Error bars are standard S.E.M.

**Supplemental Figure 6:**
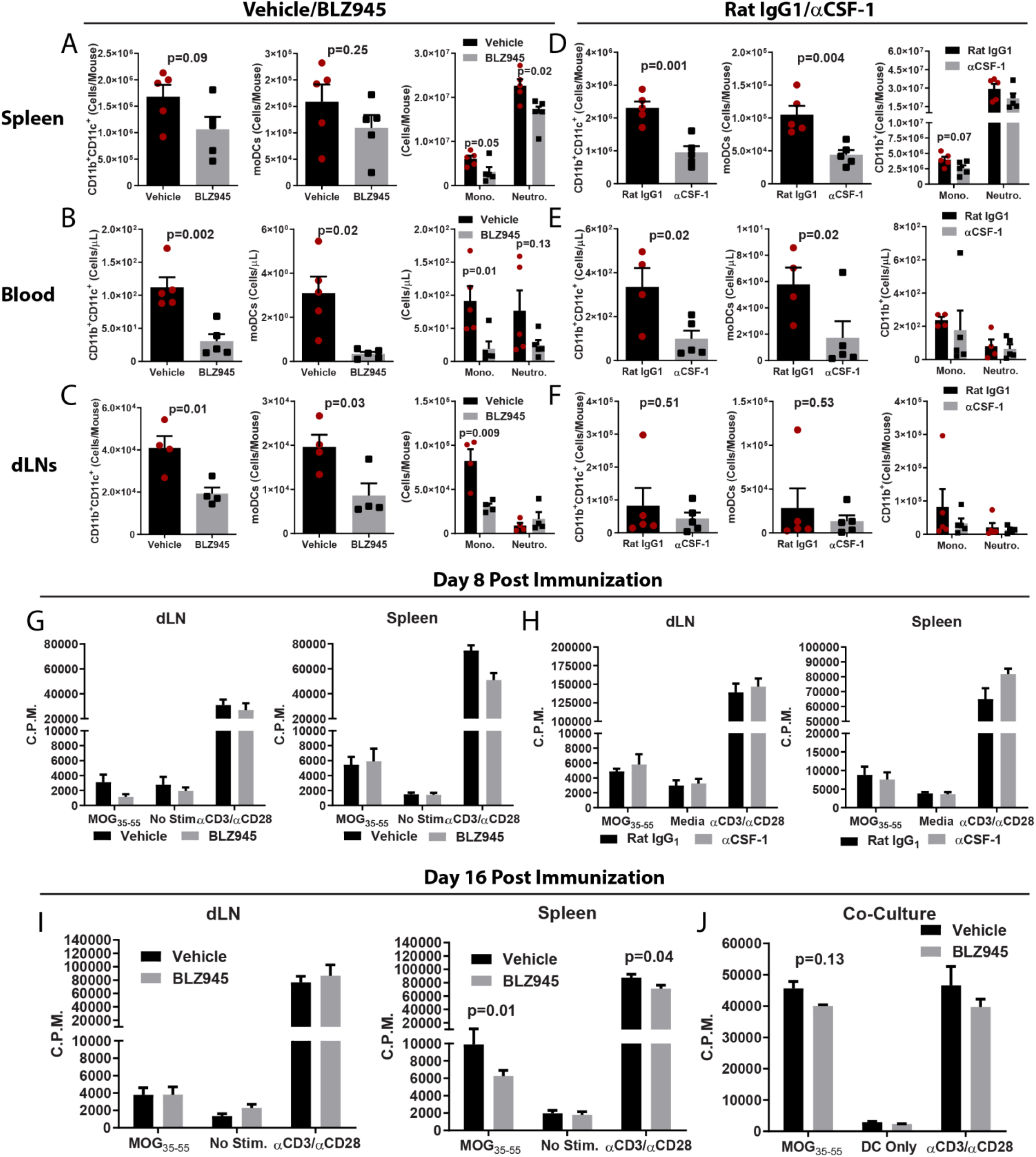
CSF-1R inhibition depletes myeloid DCs and monocytes in peripheral lymphoid compartments. Characterization of immune cells in the spleen, blood and dLNs from MOG35-55-immunized mice sacrificed on day 8 p.i.. **A-C)** BLZ945- and vehicle-treated mice (n=5 per group). Numbers of CD11b^+^CD11c^+^, moDCs, monocytes and neutrophils in **A)** spleen, **B)** blood, and **C)** dLN. **E-F)** Same quantification as in A-C) but for anti-CSF-1- and rat IgG1 control MAbs-treated mice. **G)** [^3^H]Thymidine proliferation assay for MOG35-55-stimulated splenocytes and dLNs cells from BLZ945- and vehicle-treated mice harvested on day 8 p.i. **H)** Same analyses as in G), but for anti-CSF-1- and rat IgG1 control MAb-treated mice. **I)** [^3^H]Thymidine proliferation assay for MOG35-55-stimulated splenocytes and dLNs cells from BLZ945- and vehicle-treated mice harvested on day 16 p.i. **J)** CD11c^+^ cells in splenocytes from BLZ945- and vehicle-treated mice harvested on day 8 p.i. CD11c^+^ cells were isolated by magnetic bead sorting and mixed in a 1:10 ratio with CD4^+^ T cells isolated from splenocytes of 2D2 mice, and stimulated with MOG35-55 for 72 h. Proliferation was then measured by [^3^H]Thymidine incorporation assay. Statistical significance was calculated using two-way unpaired *t* test. Error bars are S.E.M.

**Supplemental Figure 7:**
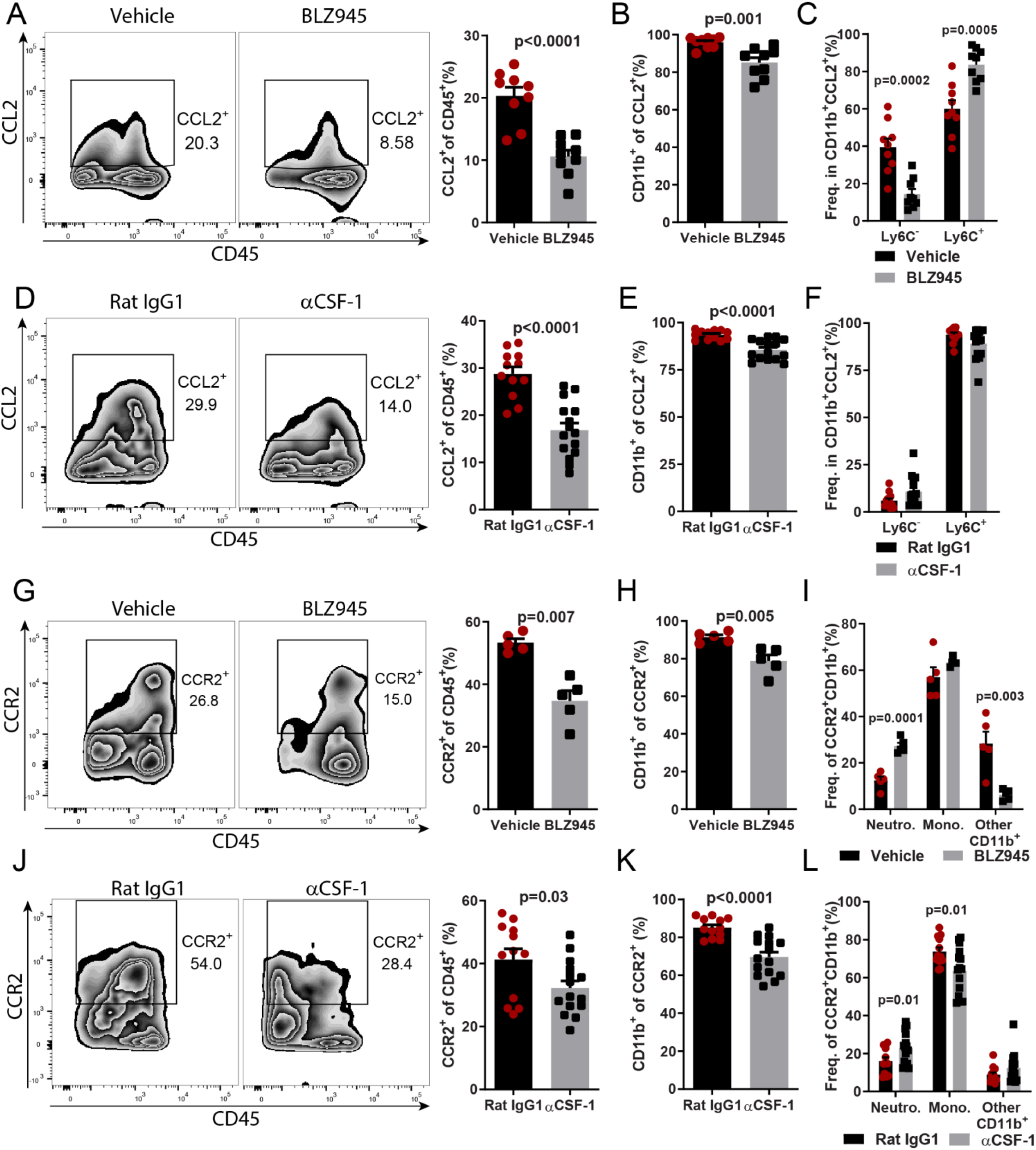
Blocking CSF-1R or CSF-1 reduces numbers of CCL2^+^ and CCR2^+^ myeloid cells in the CNS during EAE. **A)** Frequency of CCL2^+^ cells among CD45^+^ cells in the CNS of BLZ945- and vehicle-treated mice (n=9 per group) sacrificed after 6 days of treatment, at day 20 p.i.. **B)** CD11b^+^ cells among CCL2^+^ cells. **C)** Frequency of Ly6C^+^ and Ly6C^-^ cells among CD11b^+^CCL2^+^ cells. **D)** Frequency of CCL2^+^ cells among CD45^+^ cells from the CNS of anti-CSF-1- and control MAb-treated mice (n=12-15 per group), sacrificed after 6 days of treatment, at 17 days p.i.. **E)** CD11b^+^ cells among CCL2^+^ cells. **F)** Frequency of Ly6C^+^ and Ly6C^-^ cells among CD11b^+^CCL2^+^ cells. **G)** Frequency of CCR2^+^ cells among CD45^+^ cells from the CNS of BLZ945- and vehicle-treated mice from A). **H)** CD11b^+^ cells among CCL2^+^ cells. **I)** Frequency of neutrophils (CD45^Hi^CD11b^+^Ly6G^Hi^Ly6C^Int^), monocytes/monocyte-derived cells (CD45^Hi^CD11b^+^Ly6G^Neg/Lo^Ly6C^+^) and other CD11b^+^ cells among CCR2^+^CD11b^+^ cells. **J)** Frequency of CCR2^+^ cells among CD45^+^ cells from the CNS of anti-CSF-1- and control MAb-treated mice from D). **K)** CD11b^+^ cells among CCL2^+^ cells. **L)** Frequency of neutrophils (CD45^Hi^CD11b^+^Ly6G^Hi^Ly6C^Int^), monocytes/monocyte-derived cells (CD45^Hi^CD11b^+^Ly6G^Neg/Lo^Ly6C^+^) and other CD11b^+^ cells among CCR2^+^CD11b^+^ cells. Statistical significance was calculated using two-tailed unpaired *t* test. Error bars are S.E.M.

**Supplemental Figure 8.**
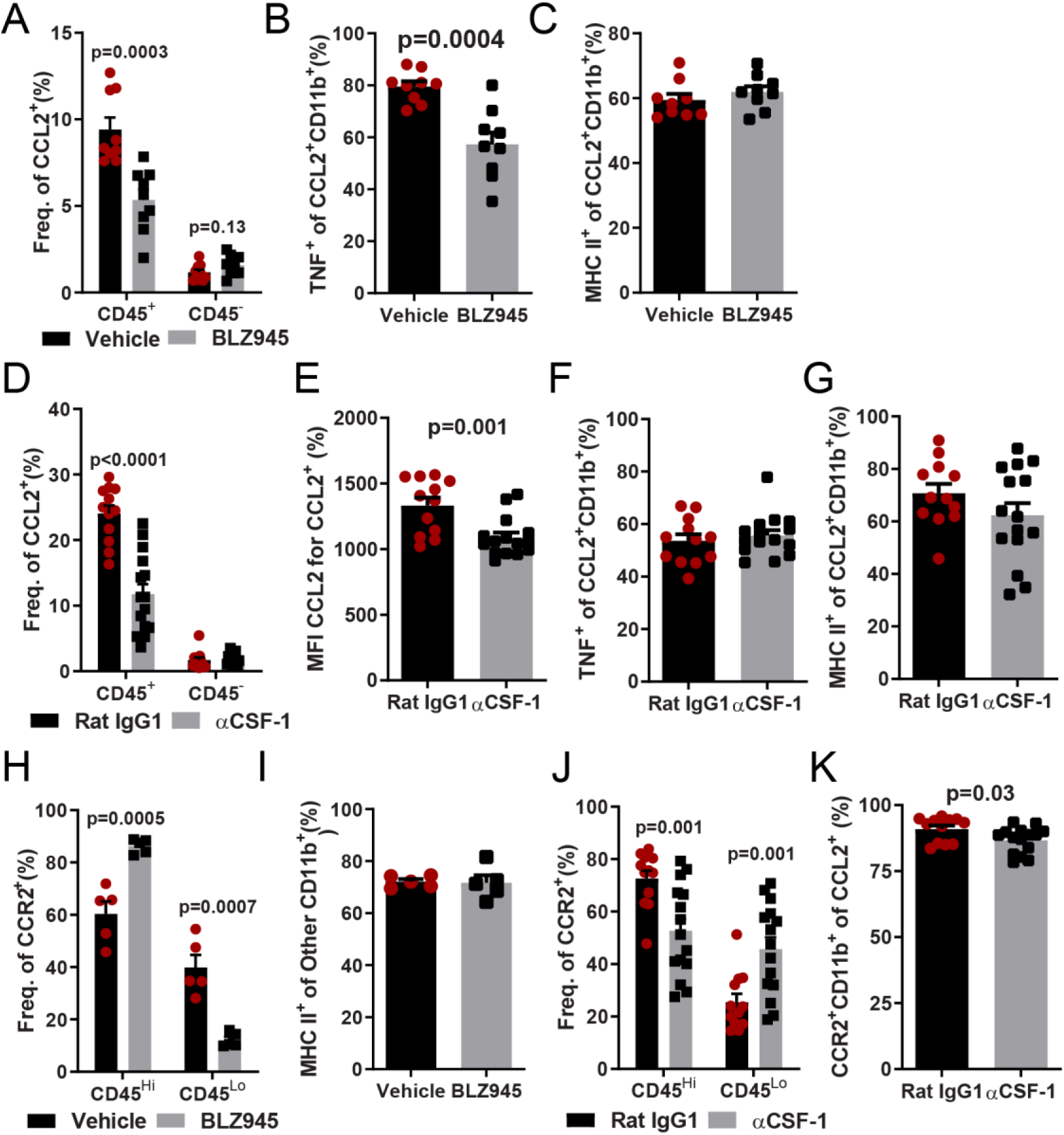
CCL2^+^ and CCR2^+^ cells are primarily composed of inflammatory myeloid cells. Frequency of CCL2^+^CD45^+^ and CCL2^+^CD45^-^ cells among mononuclear cells isolated from the CNS of BLZ945- and vehicle-treated mice with EAE. **B)** Frequency of TNF^+^, and **C)** MHC II^+^ cells among CCL2^+^CD11b^+^ cells. **D)** Frequency of CCL2^+^CD45^+^ and CCL2^+^CD45^-^ cells among mononuclear cells isolated from the CNS of anti-CSF-1- and control MAb-treated EAE mice. **E)** MFI of CCL2^+^ cells among CD45^+^CCL2^+^ cells. **F)** Frequency of TNF^+^, and **G)** MHC II^+^ cells among CCL2^+^CD11b^+^ cells. **H)** Frequency of CCR2^+^ cells among CD45^+^ cells that were either CD45^Hi^ or. CD45^Lo^ in BLZ945- or vehicle-treated EAE mice. **I)** Percentage of MHC II^+^ among “Other CD11b^+^ cells^”^ from Fig. 7B. **J)** Frequency of CCR2^+^ cells among CD45^+^ cells that were either CD45^Hi^ or CD45^Lo^ in anti-CSF-1- or control MAb-treated mice. **K)** Frequency of CCR2^+^CD11b^+^ cells among CCL2^+^ cells. Statistical significance was calculated with two-tailed unpaired *t* test. Error bars are S.E.M.

**Supplemental Figure 9.**
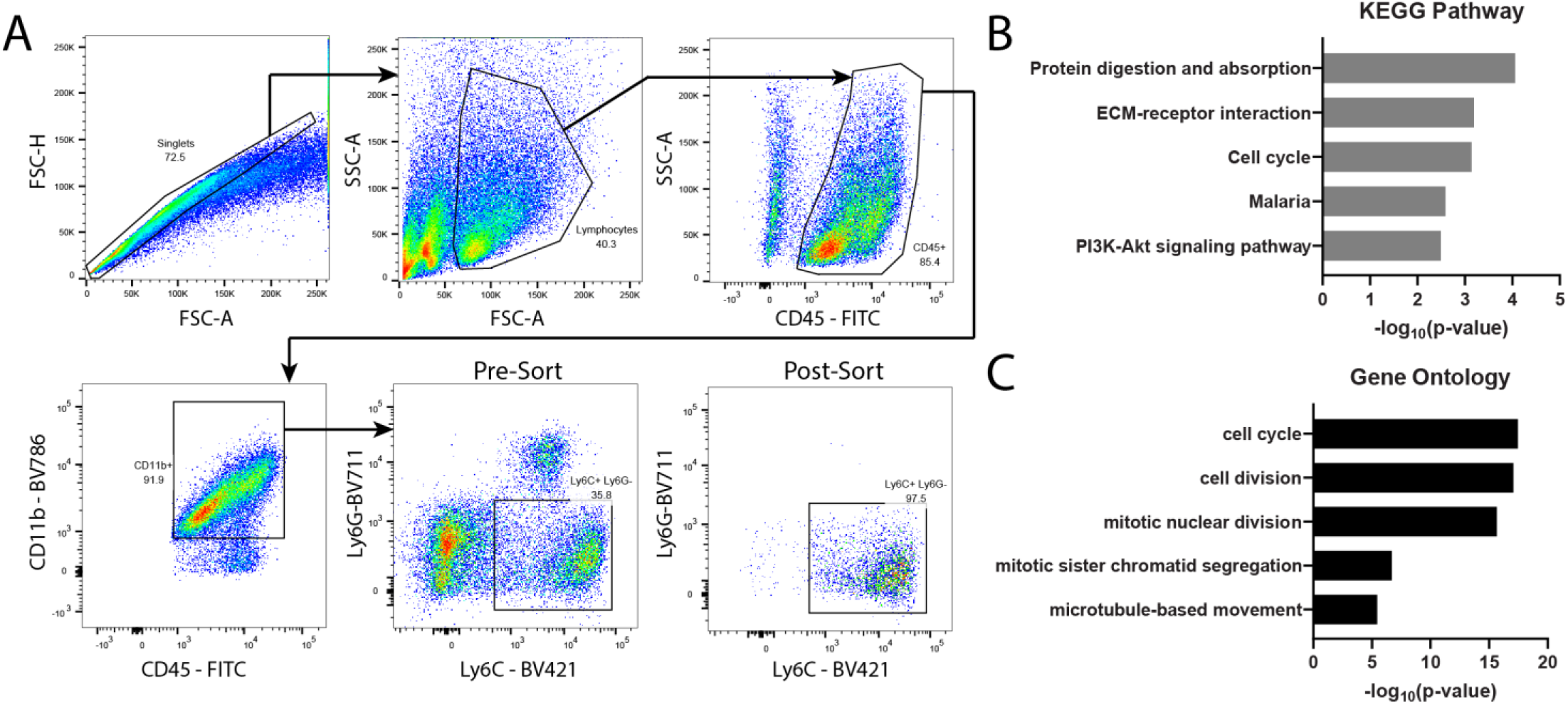
Monocytes from the CNS of mice with EAE treated with anti-CSF-1 MAb have distinct transcriptional profile. EAE was induced in C57BL/6J mice by immunization with MOG35-55. Mice were treated with 200 μg/day of anti-CSF-1 MAb or isotype control MAb from day 11 to 16 p.i. and sacrificed on day 17 p.i. **A)** Gating strategy for FACS sorting of CD45^Hi^CD11b^+^Ly6G^-^Ly6C^+^ monocytes/monocyte-derived cells. **B)** Gene ontology and **C)** KEGG pathway terms ranked by significance and generated from differentially expressed genes between monocytes/monocyte derived cells from anti-CSF-1- and isotype control MAb-treated mice.

**Supplemental Figure 10.**
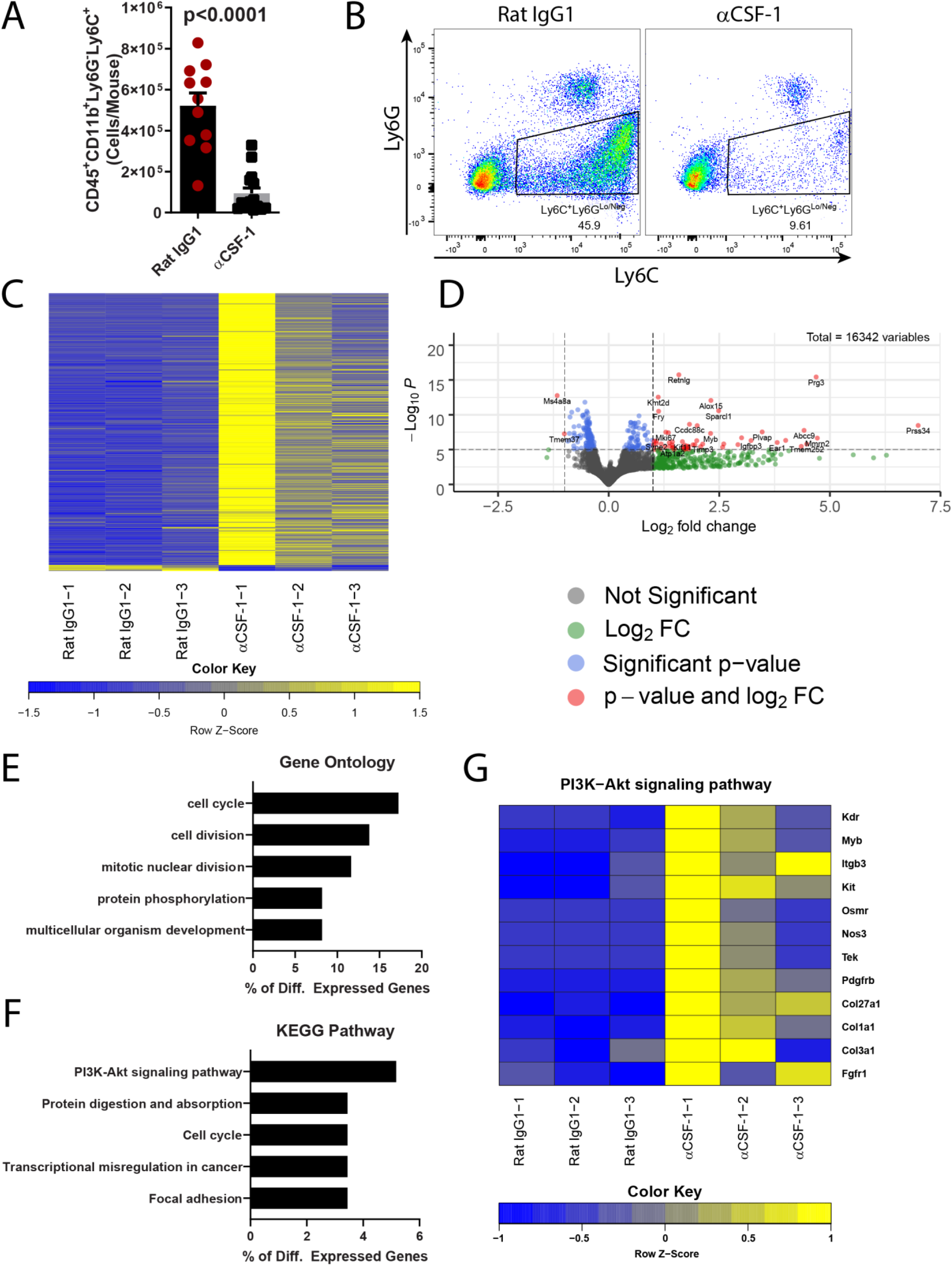
Monocytes from the CNS of anti-CSF-1 MAb-treated mice with EAE have a transcriptionally distinct phenotype. EAE was induced in C57BL/6J mice by immunization with MOG35-55. Mice were treated with 200 μg/day of anti-CSF-1 MAb or isotype IgG1 control MAb (n=3/group) from day 11 to 16 p.i. and sacrificed on day 17 p.i. **A)** Numbers of CD45^Hi^CD11b^+^Ly6G^-^Ly6C^+^ monocytes. **B)** Flow cytometry plots showing expression of Ly6C and Ly6G among CD45^+^CD11b^+^ cells. **C)** Heatmap showing differentially expressed transcripts in monocytes from anti-CSF-1- and isotype MAb-treated mice. Criteria for inclusion in heatmap were expression in all samples, with a p-adjusted value < 0.05 and a log2 fold change greater/less than ±1. **D)** Volcano plot showing relative expression of transcripts detected in samples from anti-CSF-1- and isotype MAb-treated mice. **E)** Gene ontology and **F)** KEGG pathway terms that were significantly enriched (p<0.05) and ranked by percentage of differentially expressed genes (p- adj<0.01). **G)** Heatmap of z-score- normalized TPM data from differentially expressed genes detected from the PI3K- Akt signaling pathway KEGG term.

**Supplemental Figure 11.**
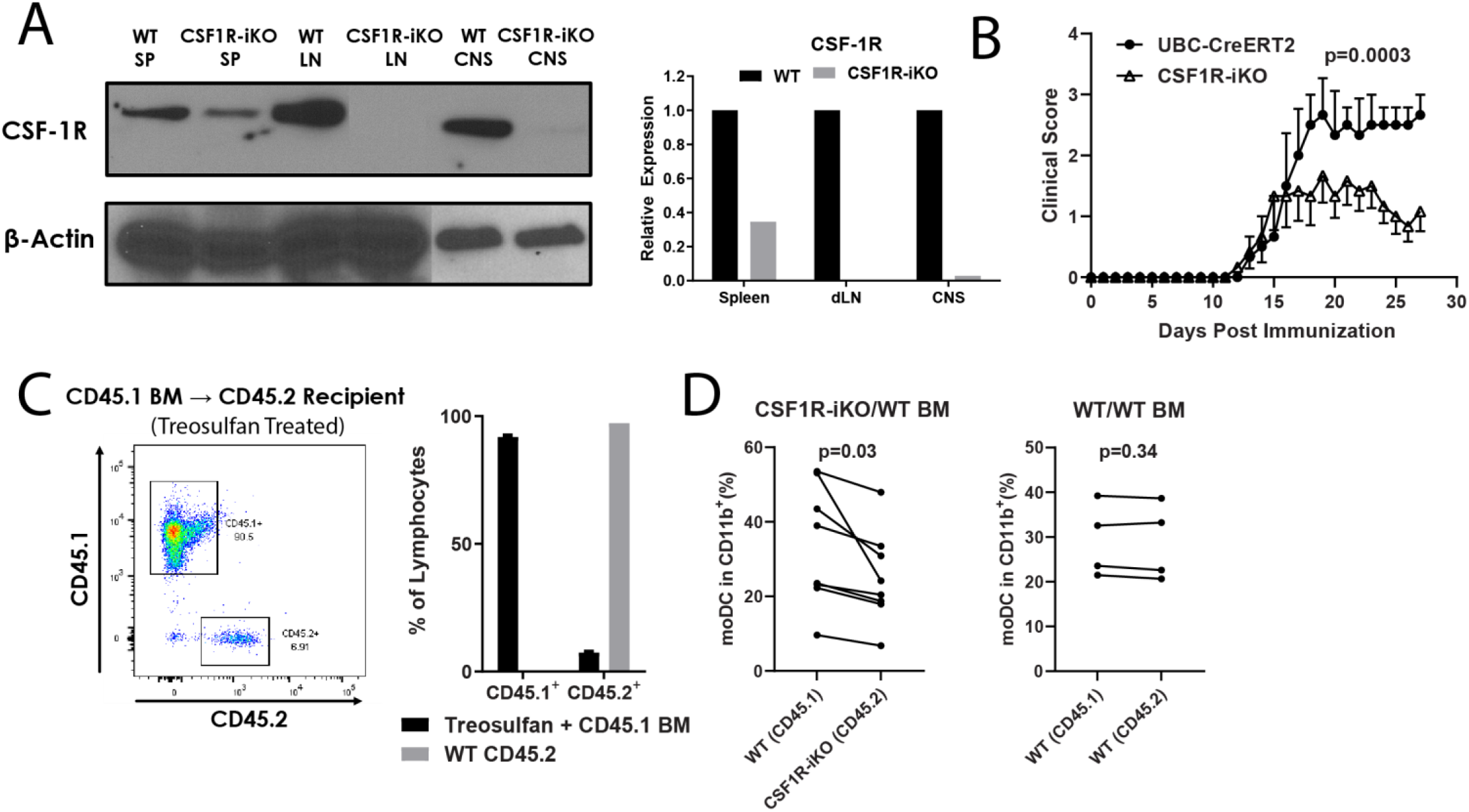
High degree of chimerism in treosulfan-induced BM chimera mice; and reduction of moDCs in CSF1R-iKO/WT mixed BM chimeras. **A)** Western blot showing protein level of CSF-1R in cell lysates of whole spleen, draining lymph nodes, and CNS mononuclear cells 5 days after 5-day treatment with tamoxifen. Relative expression, shown in the right panel, was calculated by densitometry of CSF-1R signal relative to β-actin signal, then normalized to WT expression. Each bar represents pooled lysate from 5 mice, with equal numbers of cells from each mouse pooled together. **B)** EAE in tamoxifen-treated CSF1R-iKO (n=6) and UBC-CreERT2 (n=3) mice. Significance was determined by two-way ANOVA. **C)** WT CD45.2^+^ recipient mice were treated with treosulfan and rested for 72 h, then reconstituted with 10^7^ BM cells from CD45.1^+^ mice. After 32 days, blood was collected via tail vein and reconstitution efficiency determined. Left panel shows flow cytometry plot of CD45.1 and CD45.2 expression in PBMCs. Right panel shows quantification of reconstitution efficiency. WT CD45.2 (grey bar) represents blood from naïve WT CD45.2^+^ mice. **D)** Quantification of CD45^+^Sall1^-^CD11b^+^Ly6G^-^CD11c^+^Ly6C^+^MHCII^+^ moDCs in mixed BM chimeras. Left panel shows moDCs in the CNS from mice reconstituted with a 1:1 mixture of WT and CSF1R-iKO BM cells. Right panel shows moDCs from mice reconstituted with a 1:1 mixture of CD45.1 WT and CD45.2 WT BM cells. moDCs derived from CD45.1 and CD45.2 immune cells in the CNS were compared. Significance was determined by paired *t* test.

